# Motif-based phosphoproteome clustering improves modeling and interpretation

**DOI:** 10.1101/2021.06.09.447799

**Authors:** Marc Creixell, Aaron S. Meyer

## Abstract

Cell signaling is orchestrated in part through a network of protein kinases and phosphatases. Dysregulation of kinase signaling is widespread in diseases such as cancer and is readily targetable through inhibitors of kinase enzymatic activity. Mass spectrometry-based analysis of kinase signaling can provide a global view of kinase signaling regulation but making sense of these data is complicated by its stochastic coverage of the proteome, measurement of substrates rather than kinase signaling itself, and the scale of the data collected. Here, we implement a dual data and motif clustering strategy (DDMC) that simultaneously clusters substrate peptides into similarly regulated groups based on their variation within an experiment and their sequence profile. We show that this can help to identify putative upstream kinases and supply more robust clustering. We apply this clustering to large-scale clinical proteomic profiling of lung cancer and identify conserved proteomic signatures of tumorigenicity, genetic mutations, and tumor immune infiltration. We propose that DDMC provides a general and flexible clustering strategy for the analysis of phosphoproteomic data.

**One-sentence Summary:** DDMC is a general and flexible strategy for phosphoproteomic analysis by clustering phosphopeptides using both their phosphorylation abundance and sequence motifs.

## Introduction

Cell signaling networks formed by protein kinases dictate cell fate and behavior through protein phosphorylation (*1*). As such, it is not surprising that kinase dysregulation orchestrates the onset and development of a myriad of diseases, including cancer. Measuring cell signaling by mass spectrometry (MS)-based global phosphoproteomics provides a promising opportunity to direct therapy development (*2*), particularly given the accessibility of these signaling changes to drug targeting. Nevertheless, despite the rapid accumulation of large-scale phosphoproteomic clinical data, it is still difficult to identify the signaling events leading to observed proteomic alterations and phenotypic outcomes.

One approach to make sense of phosphoproteomic measurements has been to infer the activity of upstream kinases. Previously published methods have combined each phosphopeptide with reported kinase-substrate interactions to reconstruct signaling networks. For instance, kinase-substrate enrichment analysis (KSEA) averages the signals of groups of kinase substrates to infer enriched pathways in biological samples (*3*). Another method, Integrative Inferred Kinase Activity (INKA), infers kinase activity by integrating the scores of two components that compute kinase’s overall and activation loop phosphorylation alongside another two components that quantify the phosphorylation abundance of known substrates. Kinase-substrate relationships are either experimentally determined or predicted by NetworKIN, an algorithm that uses sequence motif and protein-protein network information (*4–6*). Finally, Scansite predicts kinase-substrate interactions using sequence motifs generated from oriented peptide library scanning experiments (*7*). These methods, sometimes in combination, help to reconstruct signaling pathway activities from phosphoproteomic measurements.

Kinase-substrate inference still provides a limited view of signaling network changes, however. Kinase prediction methods are necessarily dependent on having well-characterized kinase-substrate interactions. Unfortunately, the majority of the phosphoproteome remains largely uncharacterized (*8*). Just 20% of kinases have been shown to phosphorylate 87% of currently annotated substrates and around 80% of kinases have fewer than 20 substrates, with 30% yet to be assigned a single substrate (*8*). Hence, insights generated by computational methods dependent on this unequal knowledge distribution are less likely to identify understudied protein kinases. An additional major challenge being faced during the analysis of large-scale signaling data is missingness. This is due to two major limitations of discovery-mode multiplexed tandem mass tag (TMT) MS. The technique processes batches of samples with stochastic signaling coverage in each experiment. This means that the portion of the phosphoproteome quantified in the samples of different TMT experiments varies (*9*). Thus, in the resulting data set, phosphosites are observed in certain groups of samples but not others. Computational tools usually require complete data sets and so a frequent strategy to handle this challenge is either imputing missing values with a representative statistic (e.g. average signal) or throwing out any peptides displaying missing values–at the expense of losing critical information (*10, 11*). Kinase enrichment and prediction methods are further compromised by this problem. Thus, there is a clear need to develop tailored and unbiased computational methods capable of modeling the entirety of the phosphoproteomic data set despite missingness.

Clustering methods such as hierarchical or k-means clustering identify signaling nodes by grouping phosphopeptides based on their co-variation. This clustering criterion results in groups of peptides that display similar activation patterns across conditions, but that may be targeted by sets of different upstream kinases. The residues surrounding phosphorylation sites have had to evolve throughout millions of years to become exquisitely fine-tuned motifs that confer signaling specificity and fidelity (*12, 13*). Clustering based on motif similarity might, therefore, improve model interpretation by facilitating the identification of upstream kinases modulating particular clusters that display conserved sequence motifs. On the other hand, clustering peptides based on sequence distance may result in groups of proteins that, while sharing the same set of upstream kinases, are differently regulated due to context. Thus, combining phosphorylation status and sequence similarity may enable a balanced characterization of the cell signaling state.

Here, we present an algorithm, Dual Data and Motif Clustering (DDMC), that probabilistically and simultaneously models both the peptide phosphorylation variation and peptide sequence motifs of peptide clusters to reconstitute cell signaling networks and identify causal interactions (Fig. 1). To test the utility of our method, we analyze the phosphoproteomes of 110 treatment-naïve lung adenocarcinoma (LUAD) tumors and 101 paired normal adjacent tissues (NATs) from the National Cancer Institute (NCI)’s Clinical Proteomic Tumor Analysis Consortium (CPTAC) LUAD study (*11*). We characterize the phosphoproteome of patients by identifying those signaling signatures associated with tumorigenesis, the presence of specific mutations, and tumor immune infiltration. In total, we demonstrate DDMC as a general strategy for improving the analysis of phosphoproteomic surveys.

**Figure 1:**
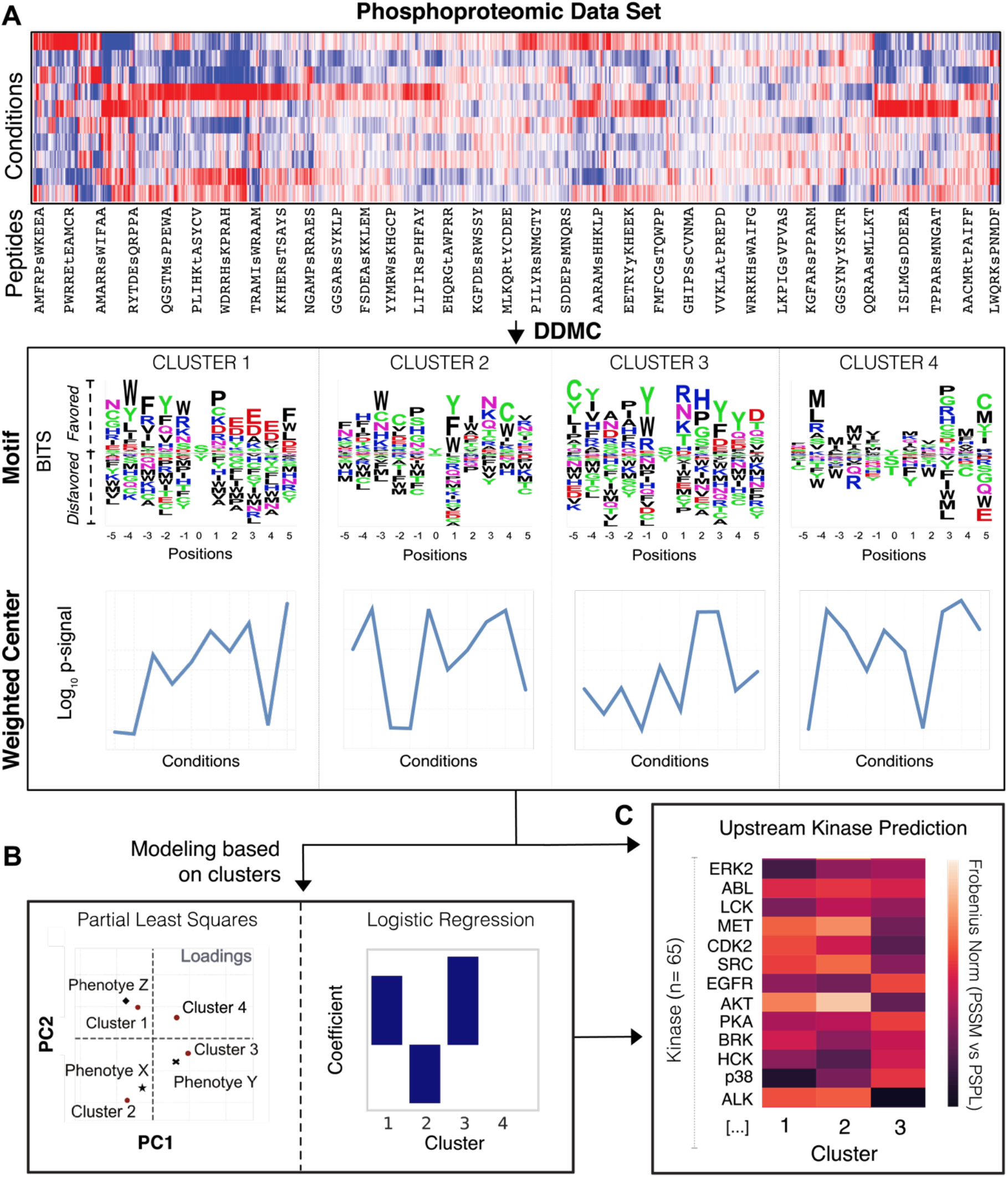
Schematic of the DDMC approach to cluster global signaling data and infer upstream kinases driving phenotypes. A) DDMC is run to cluster an input phosphoproteomic data set to generate 4 clusters of peptides that show similar sequence motifs and phosphorylation behavior. B) Predictive modeling using clusters allows one to establish associations between specific clusters and features of interest. C) Putative upstream kinases regulating meaningful clusters can be predicted by computing the distance between a cluster motif and PSPL PSSM. PSSM; Position-specific scoring matrix, PSPL; Position scanning peptide library.

## Results

### Constructing an expectation-maximization algorithm tailored for clustering phosphoproteomic data

MS-based global phosphoproteomic data provides unparalleled coverage when interrogating kinase signaling networks and their therapeutic implications. However, these data also present challenging issues as a consequence of their incomplete and stochastic coverage, high-content but low-sample throughput, and variation in coverage across experiments. In addressing these issues, we recognized that MS measurements provide two pieces of information: the exact site of phosphorylation on a peptide sequence and some measure of abundance within the measured samples. Both of these pieces of information are critical to the overall interpretation of the data.

Based on this observation, we built a mixture model that probabilistically clusters phosphosites based on both their peptide sequence and abundance across samples (Figure S1). In each iteration, DDMC applies an expectation-maximization algorithm to optimize clusters that capture the average features of member sequences and their abundance variation (Figure 1A and S1). Both information sources—peptide abundance and sequence—can be prioritized by a weight parameter. With a weight of 0, DDMC becomes a Gaussian Mixture Model (GMM) that clusters peptides according to their phosphorylation signal. With a very large weight, DDMC exclusively clusters peptides according to their peptide sequences. Clustering both the sequence and abundance measurements ensures that the resulting clusters are a function of both features, which we hypothesized would provide both more meaningful and robust clusters.

The resulting clustering provides coordinated outputs that can be used in a few different ways. The cluster centers, by virtue of being a summary for the abundance changes of these peptides, can be regressed against phenotypic responses (e.g., cell phenotypes or clinical outcomes) to establish associations between particular clusters and response (Figure 1B). Regression using the clusters instead of each peptide ensures that the model can be developed despite relatively few samples, with minimal loss of information since each peptide within a cluster varies in a similar manner.

In parallel or independently, one can interrogate the resulting Position-Specific Scoring Matrices (PSSMs) to describe the overall sequence features of that cluster. These outputs can be readily compared to other information such as experimentally generated profiles of putative upstream kinases via Position Specific Scanning Libraries (PSPL) (*14-18*). We extracted a collection of 62 kinase specificity profiles to identify which cluster motifs most resemble the optimal motif of putative upstream kinases (Figure 1C) (*17-19*). However, as kinase-substrate specificity is also dictated by features outside of the immediate substrate region, we also note that our approach is more general than strictly assembling kinase-substrate predictions as non-enzymatic specificity information may be present in the DDMC sequence motifs. Overall, this overview demonstrates how DDMC can take complex, coordinated signaling measurements and find patterns in the phosphorylation signals to reconstruct signaling networks and associate particular clusters and phenotypes.

### Dual data-motif clustering strategy robustly imputes missing values

A major limitation of multiplexed MS-based large-scale phosphoproteomic data is the presence of missing values due to (i) the limited number of samples processed at a time per TMT experiment and (ii) the stochastic signaling coverage in each experiment. Consequently, upon concatenation of the different TMT experiments, many phosphosites are observed in groups of samples. To evaluate the robustness of our combined dual data-motif clustering (DDMC) method in analyzing incomplete data sets, we designed a computational experiment wherein we removed specific observations and predicted them using the cluster centers corresponding to the peptides those missing values belonged to (Figure 2A). The resulting mean squared errors between the actual and predicted values were compared to commonly used imputation strategies such as the peptides’ mean or minimum signal, constant zero, or matrix completion by PCA. Furthermore, we evaluated the imputation performance of our method when clustering the data using a different number of clusters. We observed that increasing the number of clusters improved the imputation of missing values (Figure 2B-F). Additionally, we performed the same experiment by clustering the data with different weights. Weight changes barely affected imputation performance, indicating that cluster centers based on sequence only imputed missing values as accurately as when using the phosphorylation signal (Figure 2F-I). These results indicate that DDMC clearly outperforms standard imputation strategies such as using constant zero or the peptides’ mean or minimum signal and imputes missing values with similar accuracy to matrix completion by PCA.

**Figure 2:**
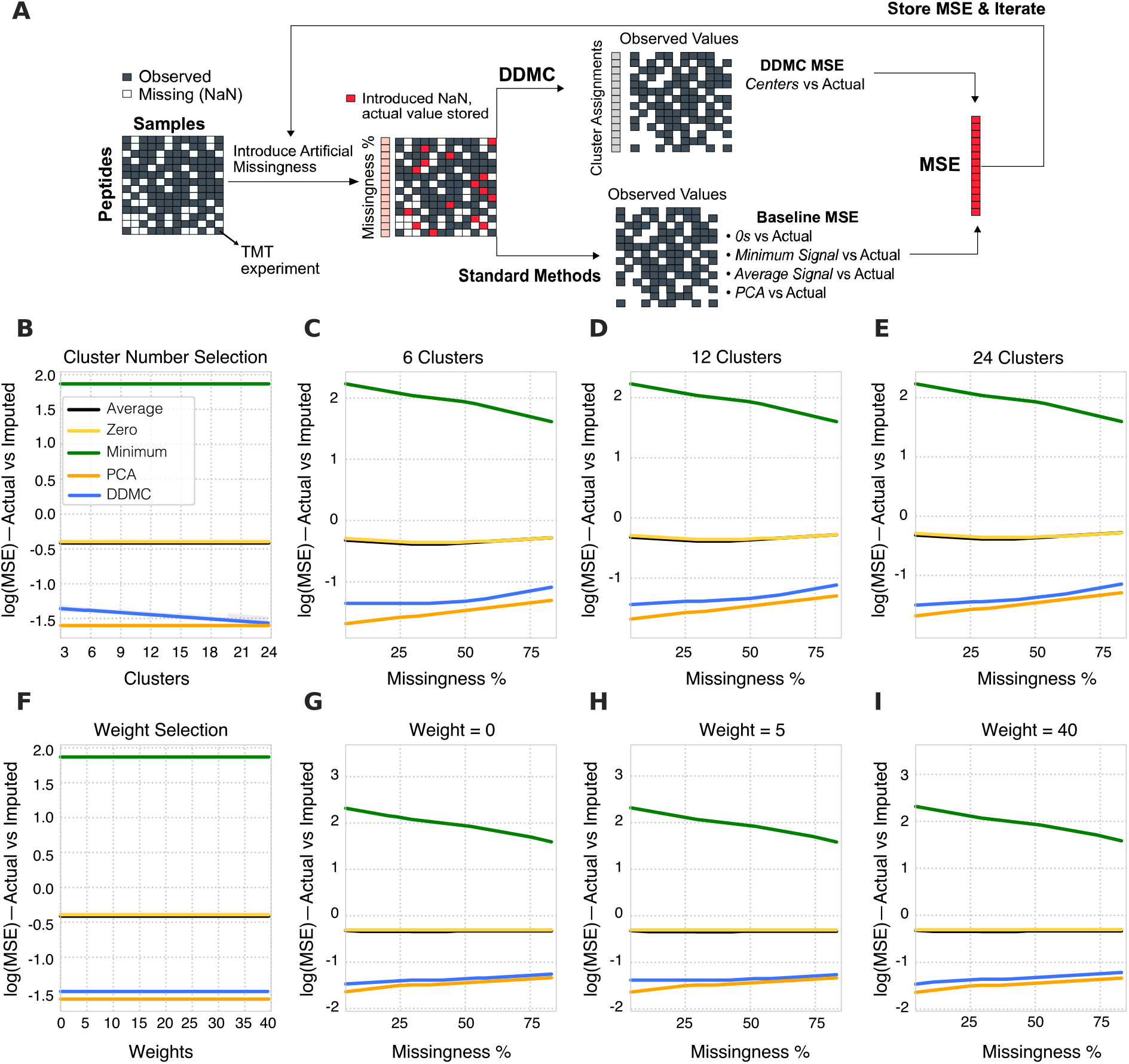
Benchmarking the robustness of motif clustering to missing measurements. A) A schematic of the process for quantifying robustness to missing values. Any peptides containing less than 7 TMT experiments were discarded. For the remaining 15904 peptides, an entire random TMT experiment was removed per peptide and these values were stored for later comparison. Next, these artificial missing values were imputed using either a baseline strategy (peptide mean/minimum signal, constant zero, or matrix completion by PCA) or the corresponding cluster center. Once a mean squared error was computed for each peptide, the second iteration repeats this process by removing a second TMT experiment. A total of 5 random TMT experiments per peptide were imputed by clustering using a different number of clusters (B-E) or different weights (E-I).

### DDMC correctly identifies AKT1 and ERK2 as upstream kinases of signaling clusters containing their substrates

DDMC is a tailored method that clusters MS-generated phosphosites using its phosphorylation behavior and sequence information. A major benefit of modeling the sequence information is the construction of cluster motifs which can be useful to infer what putative upstream kinases might preferentially target peptides of a specific cluster. To validate its ability to make upstream kinase predictions, we used DDMC to cluster the phosphoproteomic measurements of MCF7 cells treated with a panel of 61 drug inhibitors reported by Hijazi et al (*20*). PCA analysis of the resulting cluster centers clearly identified an inverse correlation between the scores of AKT/mTOR targeted inhibitors and the loading of cluster 1, indicating that the cluster’s overall signal is attenuated by the presence of these compounds (Figure 3A-B). Additional inhibitors targeting PDK1, FLT3, and S6K were also negatively correlated with cluster 1. While these do not directly inhibit AKT1/mTOR, they are all known regulators of the pathway. A heatmap displaying cluster’s 1 phosphorylation signal across treatments corroborates that the abundance of these peptides is substantially decreased when treated with AKT/mTOR/PIK3 inhibitors (Figure 3C). Encouragingly, the specificity profile of AKT—within a collection of 55 different kinase PSPL matrices—most closely matches the PSSM of cluster 1 (Figure 3D). Additionally, NetPhorest identified AKT as the second top scoring upstream kinase of cluster 1, further corroborating DDMC’s prediction.

**Figure 3:**
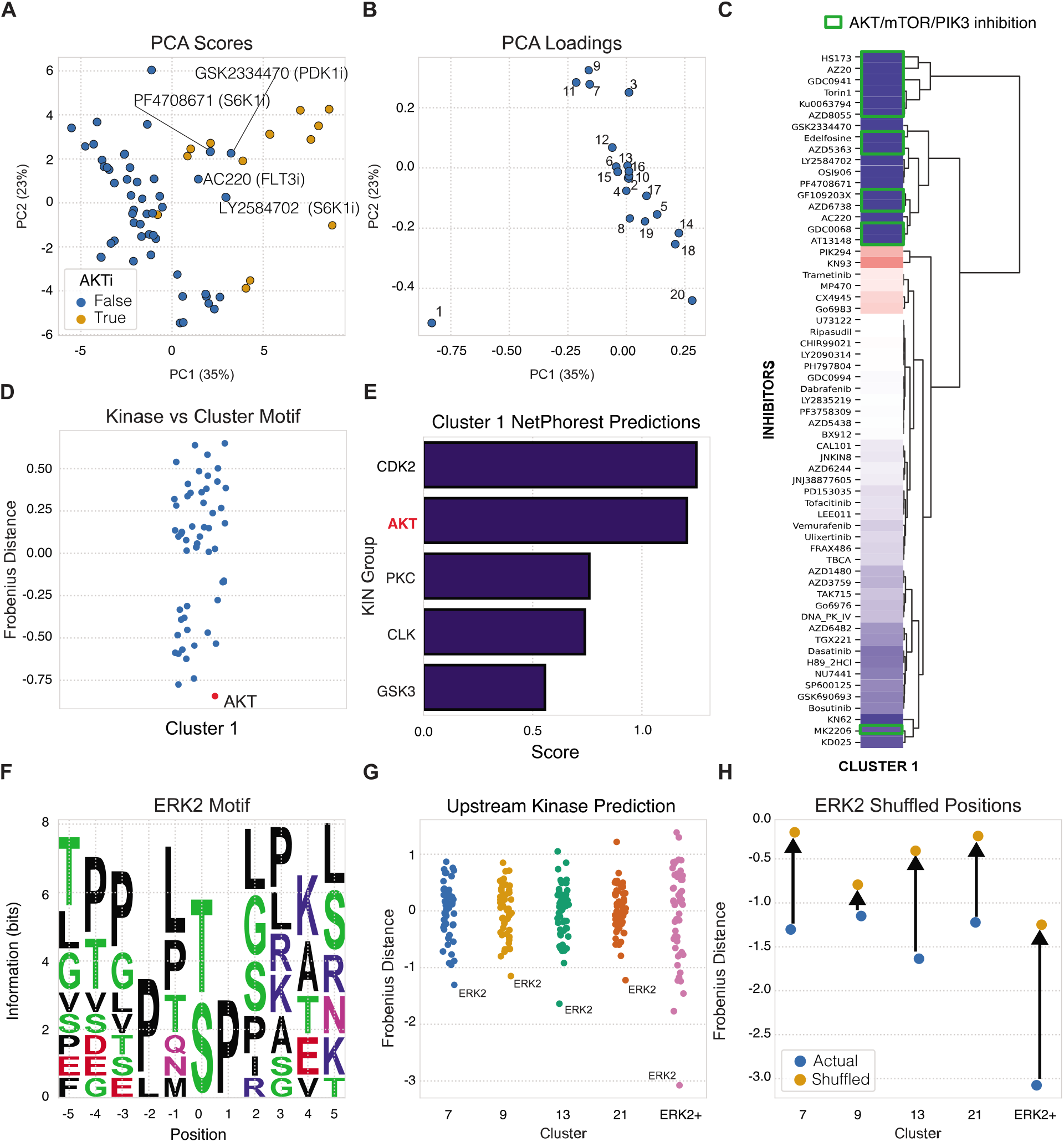
Validation of upstream kinase predictions. (A-B) PCA analysis of the DDMC phosphoproteome clusters of MCF7 cells subjected to a drug screen (20). C) Heatmap showing the effect of inhibitors on the phosphorylation signal of cluster 1. D) DDMC upstream kinase prediction of cluster 1. E) NetPhorest upstream kinase prediction of cluster 1. (F) Resulting PSSM generated using ERK2 substrates reported by Carlson et al (21). (G) Upstream kinase predictions of CPTAC clusters 7, 9, 13, and 21 in addition to the ERK2 motif shown in (F). H) Upstream kinase predictions of the same PSSMs after randomly shuffling the motif positions.

Next, we extracted the sequences of ERK2 substrates identified in Carlson *et al* to create an “artificial” ERK2-specific PSSM positive control (ERK2+ motif) (Figure 3F). As expected, ERK2 was predicted to be the upstream kinase with the highest preference for the cluster’s motif (Figure 3G). As an additional test, given the consistent enrichment of hydrophobic and polar residues throughout the entire ERK2 target motif (Figure 3F), we asked whether randomly shuffling all cluster PSSM positions surrounding the phosphoacceptor residue would affect the upstream kinase prediction. This experiment led to a 2-fold increase in the distance between ERK2 specificity profile and the ERK2+ motif (Figures 3G and H). We subjected those clusters from the CPTAC data set that were preferentially favored by ERK2 to the same experiment. As expected, we observed a similar decline in specificity between the clusters PSSMs and ERK2 PSPL matrix (Figures 3H). Note that the noticeable difference in prediction between the ERK2+ motif and the CTPAC ERK2 motifs is not surprising given that while the former group contains only 26 peptides, the CPTAC clusters contain ~500–2000 phosphosites. Overall, this experiment generally shows that despite the homogenous biophysical properties of ERK2 target motif across positions, the relative enrichment of hydrophobic and polar residues in each position determines the extent to which ERK2 favors a particular motif (Figures 3G and H). Altogether, these results illustrate two different validation scenarios in which DDMC successfully identifies the upstream kinases regulating clusters.

### A dual data-motif strategy improves prediction of different phenotypes and provides more robust clustering

As shown later in this study (Figures 5, 6, 7), we utilized DDMC to analyze the phosphoproteomes of 110 treatment-naïve LUAD tumors and 101 paired normal adjacent tissues (NATs) from the NCI’s CPTAC LUAD study. We used DDMC with the binomial sequence distance method and 24 clusters (Figure 1, 2B). We were able to include 30,561 peptides that were not observed in every tumor through our ability to handle missing data, but still filtered out 11,822 peptides that were only captured in one 10-plex TMT run. We used this fitting result throughout the rest of this study. The resulting 24 cluster motifs can be found in Figure S2.

To evaluate the benefit of incorporating the peptide sequence information into the clustering criterion, we asked whether utilizing DDMC with different sequence weights would affect the performance of a regularized logistic regression model that predicts the mutational status of STK11, whether a patient harbors a mutation in EGFR and/or a gene fusion in ALK (EGFRm/ALKf), and the level of tumor infiltration (“Hot” versus “Cold”). We found that for all three phenotypes, when the method only uses the phosphorylation signal (weight=0), the patient samples are classified with lesser accuracy compared with when a combination of both data and sequence is used. In the case of STK11, the use of the largest weight wherein mainly the sequence motifs are used for clustering provided the best prediction performance. Likewise, EGFRm/ALKf samples were best classified with a mix weight of 15 or 50. Finally, the regression model classifying whether a sample is “hot-tumor-enriched” (HTE) or “cold-tumor-enriched” (CTE) showed the best fitness with a weights of 10, 35, and 40. Together, these results indicate that observing the motif information during clustering leads to final clusters that enhance the performance of downstream phenotype prediction models (Figures 4A and S3).

**Figure 4:**
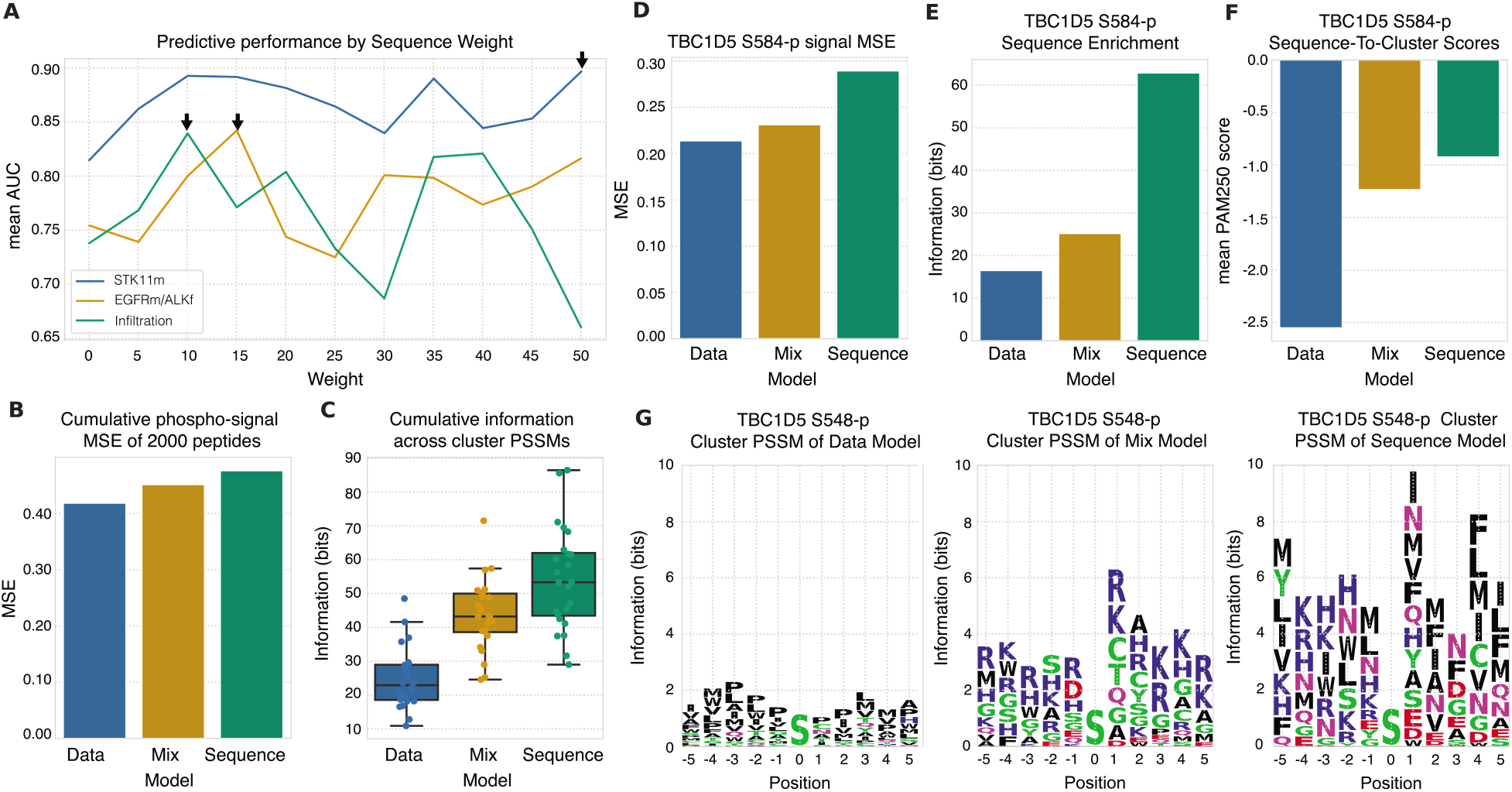
Sequence information enhances model prediction and provides more robust clustering. A) Performance of a regression model predicting the mutational status of STK11 (blue) EGFR and/or ALK (yellow) and tumor infiltration (green) in LUAD patients using either only phosphorylation data (weight=0), mainly sequence information (50), or both (0 < w < 50). B) MSE between the phosphorylation signal of 2000 randomly selected peptides and the center of its assigned clusters using a weight of 0 (data), 20 (mix), or 50 (sequence). C) Cumulative PSSM enrichment across positions comparing the data, mix, and sequence clustering strategies. (D-H) TBC1D5 peptide p-signal MSE (D), cumulative PSSM enrichment (E), and PSSM logo plots (F-H).

Next, we explored how using different weights affects the overall phosphorylation signal and sequence information of the resulting clusters. To do so, we compared the model behavior after clustering the CPTAC data with a weight of 0 (peptide abundance only), 20 (mix), and 50 (mainly sequence). First, we hypothesized that the abundance-only model would generate clusters wherein its members would show less variation in phosphorylation signal and thus a lower mean squared error (MSE). To test this, we computed the average peptide-to-cluster MSE of 2000 randomly selected peptides for each model across all clusters. Although the differences were not significant, we did observe a direct correlation between weight and MSE (Figure 4B). Next, we calculated the cumulative PSSM enrichment by summing the sequence information (bits) of all cluster PSSMs per model. As expected, increasing the weight led to a corresponding increase in the cumulative sequence information (Figure 4C). To further illustrate the clustering behavior, we tracked the phosphosite TBC1D5 S584-p in the three models. Consistent with the general trend, the abundance-only and mixed models generated lower p-signal MSE when compared to its cluster center than the Sequence model whereas weight correlated with the total PSSM enrichment (Figures 4D-E). Next, we quantified whether in addition to an increase in absolute enrichment, the mixed and sequence-only models generated more similar cluster motifs to TBC1D5 S584-p sequence than the abundance-only model. To do so, we computed the mean of all pairwise PAM250 scores between the query sequence and all cluster sequences across models which clearly confirmed that as the sequence prioritization of the model increases, the cluster PSSM is not only more enriched across all positions but also displays a more representative sequence of TBC1D5 phosphosite (Figures 4F-I). These results show that using a mixed weight that similarly prioritizes both information sources—peptide abundance and sequence—leads to more robust clustering of phosphosites through a tradeoff between phosphorylation abundance and sequence motifs.

### Widespread, dramatic signaling differences exist between tumor and normal adjacent tissue

We explored whether DDMC could recognize conserved signaling patterns in tumors compared to normal adjacent tissue (NAT). The signaling difference between tumors and NAT samples was substantial, highlighting the significant signaling rewiring that tumor cells must undergo (Figure 5A). Using principal components analysis, we could observe that NAT samples were more similar to one another than to each tumor sample (Figure 5B/C). Nearly every cluster was significantly different in its average abundance between tumor and NAT (Figure 5D). Not surprisingly given these enormous differences, samples could be almost perfectly classified using their phosphopeptide signatures, with or without DDMC (Figures 5E; S4). Using the DDMC clusters, a logistic regression model identified that NAT versus tumor status could be predicted with cluster 11 alone (Figure 5C).

**Figure 5:**
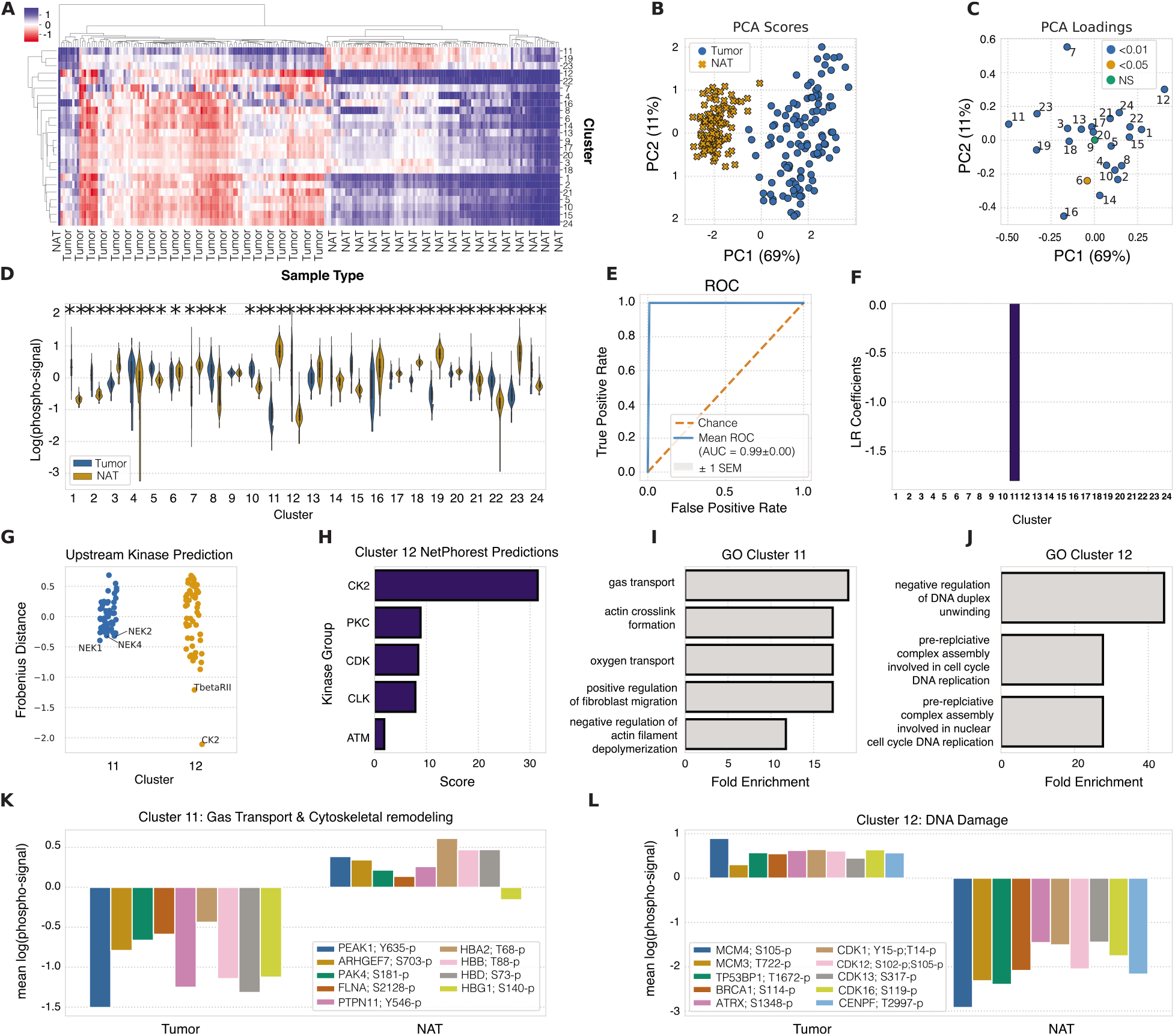
Conserved tumor differences compared to normal adjacent tissue. A) Hierarchical clustering of DDMC cluster centers. B–C) Principal components analysis scores (B) and loadings (C) of the samples and phosphopeptide clusters, respectively. D) Phosphorylation signal of tumor and NAT samples per cluster and statistical significance according to a Mann Whitney rank test (* = p-value < 0.05 and ** = p-value < 0.001). E) Receiver operating characteristic curve (ROC) of a regularized logistic regression model. F) Logistic regression weights per cluster. G) Upstream kinase predictions of clusters 11 and 12. (H) NetPhorest kinase predictions of cluster 12. (I-J) Gene ontology analysis and (K-L) representative peptides of enriched biological processes of clusters 11 and 12.

With the abundance changes and regression results we observed, we decided to further explore clusters 11 and 12. Cluster 11 shows a PSSM motif that might correspond to NEK1, 2, and 4, and an enrichment of peptides involved in gas and oxygen transport, as well as cytoskleleton remodeling or migration-related phenotypes according to a Gene Ontology (GO) analysis (Figure 5G/I). Even though NEKs are a largely understudied family of serine/threonine kinases, NEK1/2 have an established role in the formation and disassembly of cilia and NEK4 has also been implicated in regulating microtubule dynamics and stability (*22, 23*). The primary cilium serves as a signaling hub via the local expression of cell surface receptors and signaling molecules to sense environmental stimuli and thus promote a handful of phenotypes including adaptation to hypoxia, migration, and escape from apoptosis (*24, 25*). Cancer cells typically lack cilia which could promote the emergence of these malignant phenotypes. Cluster 11 displays a striking phosphorylation decrease in tumor samples compared with NATs which could be representative of the presence or lack of NEK1/2 signaling, respectively. Within this group of peptides, there is a notable overrepresentation of hemoglobin subunits (HBG1, HBD, HBB, and HBA2) which could illustrate the different oxygenation status of NATs versus malignant tissues. Moreover, several cytoskeletal-remodeling proteins are present in cluster 11 such as PEAK1, FLNA, GAS2L2, MARCKS, PEAK1, and ARHGEF7. The abundance of all these signaling molecules is substantially decreased in tumor compared to NAT samples (Figure 5K).

On the other hand, cluster 12 was clearly identified as a CK2-like motif (Figure 5G). This association was also established by NetPhorest which identified multiple experimentally validated CK2 substrates in this cluster (Figure 5J). GO analysis of cluster 12 identified a substantial enrichment of negative regulators of DNA duplex unwinding and pre-replicative complex assembly involved in cell cycle DNA replication (Figure 5G, I-J). DNA duplex unwinding and replication are important processes that play a major role in maintaining genome stability. DNA helicases are the enzymes responsible for unwinding the DNA and thus are essential for DNA replication. As such, they have been widely associated with DNA damage response (DDR) and cancer development (*26*). CK2 has been widely implicated in modulating DNA repair signaling pathways in response to DNA damage to promote cell survival in cancer (*27–29*). In fact, a study found that the CK2 inhibitor CX-4945 blocked DDR induced by gemcitabine and cisplatin and synergizes with these compounds in ovarian cancer cell lines (*30*). Cluster 12 contains several signaling proteins related to DNA replication and genome stability such as MCM3/4, the p53 interactor TP53BP1, BRCA1, ATRX, CENPF, and CDKs whose signal is strikingly decreased in NATs and increased in tumor samples (Figure 5L). These results, therefore, suggest that CK2 might activate signaling molecules within cluster 12 involved in DNA repair pathways to induce the survival of cancer cells. Taken together, DDMC builds phosphoproteomic clusters that present signaling dysregulation common to tumors compared to NATs and identifies putative upstream kinases modulating them. These features can help to interpret phosphoproteomic results and inform the generation of hypotheses for follow up experiments.

### Genetic driver mutations are associated with more targeted phosphoproteomic rewiring

Inactivating somatic mutations in STK11 lead to increased tumorigenesis and metastasis (*31*). Thus, we aimed to identify the phosphoproteomic aberrations triggered by this genetic event. The majority of clusters were significantly altered, generally toward higher abundances with a mutation (Figure 6A). The cluster centers corresponding to each patient’s tumor and NAT samples could successfully predict the STK11 mutational status by regularized logistic regression (Figure 6B). The tumor phosphoproteomic signal of cluster 7 greatly contributed to classify mutant STK11 samples, whereas the tumor signal of 8 and 14 helped classify WT STK11 specimens. (Figure 6C). These results motivated further exploration of clusters 7 and 8 which present sequence motifs favored by ERK2, and CK1/BRCA1/PKD, respectively (Figure 6D).

**Figure 6:**
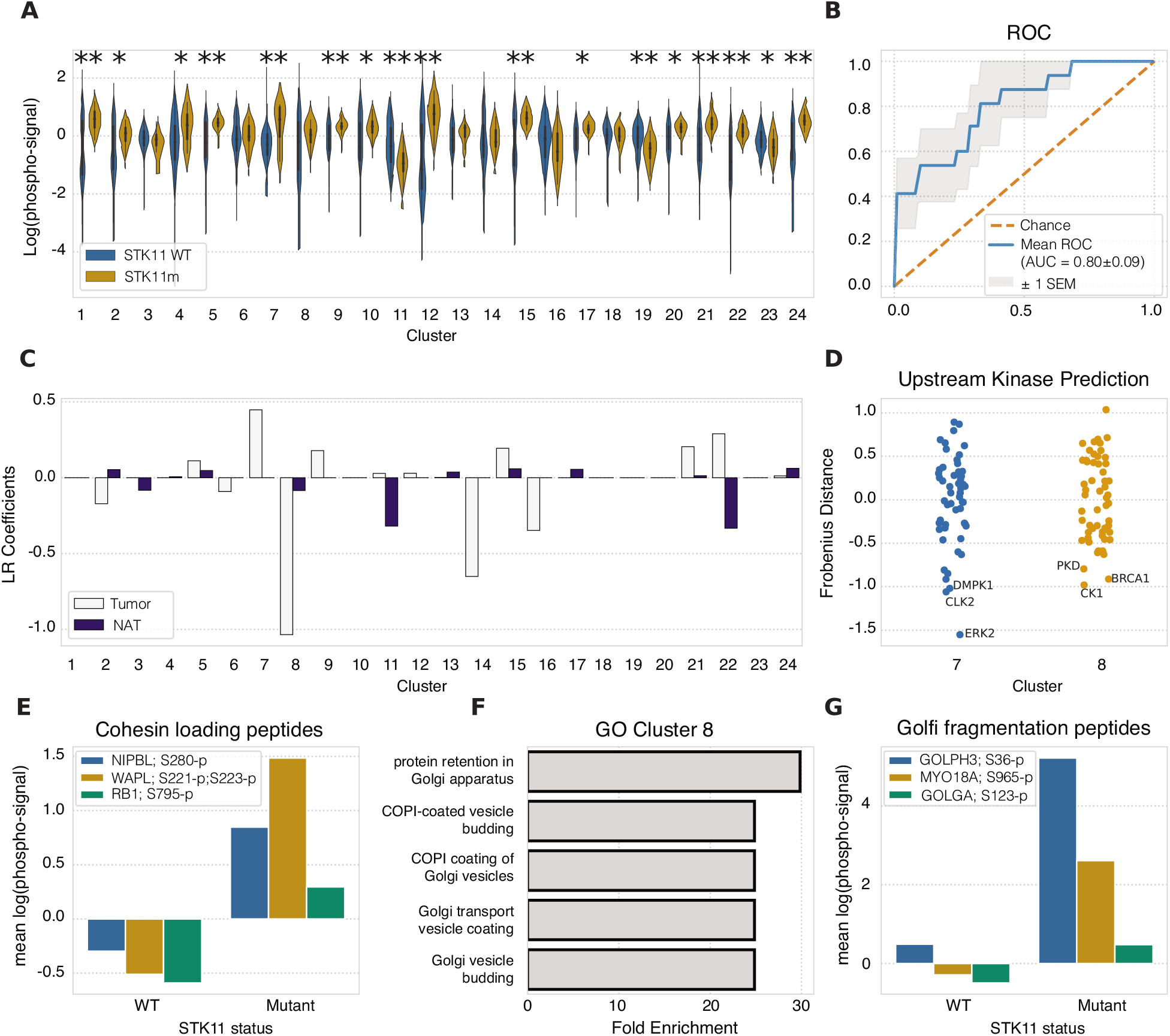
Phosphoproteomic aberrations associated with STK11 mutational status. A) Phosphorylation signal of STK11 WT and mutant samples per cluster and statistical significance according to a Mann-Whitney rank test (* = p-value < 0.05 and ** = p-value < 0.001). B) ROC of a logistic regression model predicting the STK11 mutational status and (C) its corresponding weights per sample type. (D) Putative upstream kinases of clusters 7, and 8. (E) Representative cohesin loading peptides in cluster 7. (F-G) GO analysis and representative Golgi fragmentation peptides of cluster 8.

Cluster 7 is highly enriched with peptides involved in regulation of the cell cycle by cohesin loading (Figure 6E). Cohesin is a protein complex that mediates sister chromatid cohesion by directly binding with DNA. This interaction holds both chromatids together after DNA replication until anaphase wherein cohesin is removed to facilitate chromosome segregation during cell division. Cluster 7 contains the inhibitor phosphosite of the tumor suppressor RB1 S795-p, the member of the cohesin loading complex NIPBL (S280-p, S280-p;S284-p, and S350-p), and the cohesin release factor WAPL (S221-p and S221-p;S223-p). Studies have shown that RB1 inactivation can lead to defects in chromosome cohesion that in turn compromises chromosome stability (*32, 33*). Manning et al demonstrated that depletion of WAPL in RB1-deficient cells promoted cohesin association with chromatin (*33*). Among these phosphosites, we observed strong opposing signals between STK11 WT and mutant patients in NIPBL S280-p; WAPL S221-p, S223-p; and RB1 S795-p (Figure 6E) which reinforces the association between STK11 activity and chromatin instability. Moreover, CDCA5 is key regulator of sister chromatid cohesion by stabilizing cohesin complex association with chromatin and was identified as a prognostic factor of lung cancer through a tumor tissue microarray analysis of 262 non–small cell lung cancer (NSCLC) patients (*34*). They showed that CDCA5 phosphorylation of S209 by ERK2 enhanced cell proliferation (*34*). Therefore, these results might suggest that mutations inactivating mutations in STK11 might correlate with signaling defects in sister chromatid cohesion during the cell cycle which in turn lead to chromosome instability and cell cancer growth. In fact, STK11 inactivation has been associated with genomic instability, although the signaling mechanism underlying this phenotypic response remains elusive (*35*).

The signal of phosphosites in cluster 8, specifically in tumor samples, largely contributes to predict the signaling differences between STK11 WT and mutant samples (Figure 6C). This cluster presents a clear enrichment of peptides involved in the regulation of the Golgi apparatus such as GOLGA2-5, GOLGB1, and GOLPH3 (Figure 6F). Cancer cells commonly undergo fragmentation of the Golgi which has been shown to drive several malignant molecular signatures including the hyperactivity of motor proteins and kinase signaling dysregulation (*37*). Myosin 18A and 1E pertain to cluster 18 and the former has been reported to interact with GOLPH3 to induce Golgi dispersal. Moreover, a series of studies uncovered that GOLPH3 promotes cell proliferation in cancer (*38–40*). The phosphorylation behavior of GOLPH3, Myosin 18A, and GOLGA2 in STK11 WT compared with STK11 mutant patients shows a dramatic increase of abundance in the latter which supports the association between STK11 activity and an oncogenic role of the Golgi apparatus in these patients (Figure 6E). Together, these results suggest that STK11 mutations in tumor samples could affect the dispersion of the Golgi apparatus compared with STK11 WT samples.

Tyrosine kinase inhibitors (TKIs) targeting the receptor tyrosine kinases (RTKs) EGFR and ALK are effective treatments in cancer patients with EGFR mutations and/or ALK translocations (EGFRm/ALKf). However, these treatments are limited by drug resistance which in some cases can be mediated by the concomitant signaling of both RTKs activated by driver mutations (*41, 42*). Once again, the signaling cluster centers allowed a regularized logistic regression model to more accurately classify samples according to its EGFRm/ALKf status (Figure S5).

Finally, we compared the classification performance of four regularized logistic regression models fit to either the DDMC clusters, clusters generated by the standard methods GMM and k-means, or the raw phosphoproteomic data directly. It is worth noting that unlike DDMC, methods such as GMM, k-means, or direct regression cannot handle missing values and thus for these strategies we used the 1,311 peptides that were observed in all samples, whereas DDMC was fit to the entire data set comprising 30,561 phosphosites. We found that samples were classified with higher accuracy using DDMC compared to a GMM and with similar performance to k-means, especially with STK11 (Figure S6A). Direct regression to the raw signaling data yielded excellent performance; however, this strategy assigns thousands of coefficients to different peptides that vary every time the model is run, rendering this approach unable to establish a consistent link between mutations and signaling (Figure S6). In contrast, our analysis identifies a consistent association between STK11 activity with two novel phenotypes, namely chromosome cohesion during cell cycle and Golgi fragmentation, and proposes putative signaling mechanisms to support it.

### Exploration of immune infiltration-associated signaling patterns in tumors

Immune checkpoint inhibitors (ICIs) have emerged as effective treatment options for NSCLC patients. However, there still is a need to identify or influence which patients will respond to these therapies. Patients that do not respond to ICIs often have tumors with poor immune infiltration either inherently or via an adaptive process after long exposure to the drug. However, the signaling mechanism by which malignant cells prevent tumor infiltration remains elusive. We used our DDMC clusters to explore the shared signaling patterns that differentiate “hot-tumor-enriched” (HTE) from “cold-tumor-enriched” CTE LUAD patients (*11, 43*). HTE and CTE status per patient was determined using xCell by Gilette et al (*11*).

We observed that four clusters were significantly different in their average abundance between HTE and CTE samples (Figure 7A). Cluster 17, 18, and 20 display significantly higher abundances in HTE compared to CTE samples whereas cluster 21 presents the opposite trend. Samples could be accurately classified using the DDMC clusters (Figure 7B). This predictive performance was mainly explained by a positive association of cluster 2 with HTE status and cluster 6 with CTE. Other clusters contributed to explain the signaling differences between both groups but to a lesser extent (Figure 7C).

**Figure 7:**
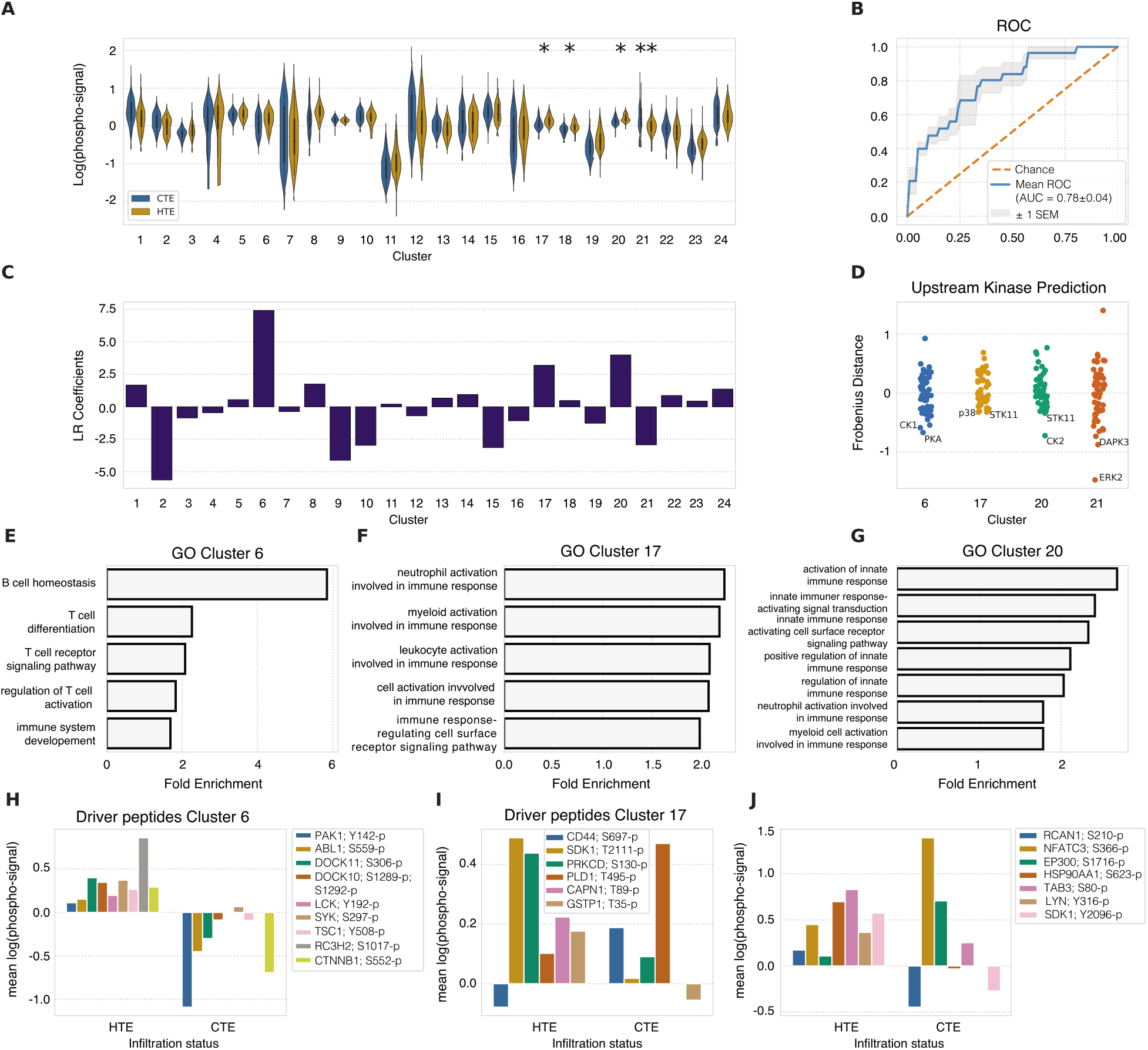
Phosphoproteomic signatures driving tumor immune infiltration. (A) Phosphorylation abundance of CTE and HTE samples per cluster and statistical significance according to a Mann-Whitney rank test (* = p-value < 0.05 and ** = p-value < 0.001). (B–C) ROC and coefficients of a logistic regression model predicting infiltration status—cold-tumor enriched (CTE) versus hot-tumor enriched (HTE). (D) Putative upstream kinases of clusters 7, 17, 20, and 21. (E–G) GO enrichment analysis of select clusters. (H–J) Selected peptides driving the GO biological processes in HTE versus CTE samples.

These results prompted us to further investigate clusters 6, 17, 20, and 21 which our model predicts to be regulated by CK1/PKA, STK11/p38, CK2/STK11, and ERK2, respectively (Figure 7D). When exploring immunologically relevant phenotypes in the GO analysis of each cluster, we observed that clusters 6, 17, and 20 showed a substantial over-representation of immunological processes. Conversely, neither of these were present in the GO analyses of cluster 2 nor cluster 21 wherein the former substantially contributes to predict CTE samples and the latter shows a significant increase of phosphorylation abundance in CTE over HTE samples (Figures 7A and C). A gene ontology analysis indicates that cluster 6 members are particularly involved in mediating B cell homeostasis, but also T cell differentiation, T cell receptor signaling, and regulation of T cell activation. These processes are promoted, at least in part, by ABL1, LCK, PAK1, and DOCK10/11 which show an increased abundance in HTE and are attenuated in CTE samples (Figures 7E & H). Cluster 17 GO analysis unveiled an over-representation of several innate and adaptive immune response pathways possibly involving CD44, SDK1, PKC, PLD1, CAPN1 and GSTP1. For instance, CD44 is expressed in both endothelial and immune cells and its regulation plays a key role in enabling neutrophil and lymphocyte recruitment into tissues (*44, 45*) (Figures 7F & I). A study found that the osteopontin (OPN)/CD44 interaction is an immune checkpoint that controls CD8+ T cell activation and tumor immune evasion in which elevated expression of OPN correlated with decreased patient survival and conferred host tumor immune tolerance. Cluster 20 is enriched in responses orchestrated by the innate immune system (Figures 7G and J). The transcription factor NFATC crucially promotes T cell activation and proliferation, and several studies show that the predicted upstream kinase of cluster 20 CK2 directly phosphorylates this protein and enhances its gene expression (*46, 47*). In addition, CK2 has also been shown to phosphorylate Regulators of Calcineurin (RCAN) proteins, which indirectly inhibit NFATC function (*48*). Several RCAN and NFATC peptides are present in cluster 20, however S210-p and S366-p, respectively show the largest abundance difference between HTE and CTE. Unexpectedly, RCAN1 S210-p shows a higher signal in HTE than in CTE whereas NFATC3 S366-p presents the opposite trend which might indicate that both phosphorylation events are inhibitory. Together, these results reinforce the role of CK2 in promoting immune infiltration in lung cancer patients. Intriguingly, inactivating mutations in STK11 have been reported to promote anti-PD1/PD-L1 resistance in KRAS-mutant LUAD suggesting a key role of STK11 in promoting tumor immune infiltration (*49*). Overall, these data demonstrate that the presence or lack of tumor immune infiltration can be accurately predicted by the DDMC clusters which in turn help identify putative upstream kinases modulating immune evasion.

## Discussion

Phosphorylation-based cell signaling through the coordinated activity of protein kinases enables cells to swiftly integrate environmental cues and orchestrate a myriad of biological processes. MS-based global phosphoproteomic data provides the unique opportunity to globally interrogate signaling networks to better understand cellular decision-making and its therapeutic implications. However, these data also present challenging issues as a consequence of their incomplete and stochastic coverage, high-content but low-sample throughput, and variation in coverage across experiments. Here, we propose a clustering method, Dual Data and Motif Clustering (DDMC), that untangles highly complex coordinated signaling changes by grouping phosphopeptides based on their phosphorylation behavior and sequence similarity (Figure 1). To test the utility of DDMC, we clustered the phosphoproteomes of LUAD patients and used the resulting groups of peptides to decipher signaling dysregulation common to tumors, genetic backgrounds, and tumor infiltration status (Figures 5, 6, 7).

Previous efforts in regressing mass spectrometry-based phosphorylation measurements against phenotypic or clinical data have been based on the ability of certain regression models such as PLSR or LASSO to robustly predict using high-dimensional and correlated data (*50*). While these models can generally be predictive with such data, they are not easily interpretable (Figure S4B). Hence, we hypothesized that clustering large-scale MS measurements based on biologically meaningful features and utilizing the cluster centers to fit regression methods could enhance the predictive performance of the model while providing highly interpretable results wherein clusters constitute signaling nodes distinctly correlated with cell patient phenotypes. Here, we demonstrate that DDMC enhances model prediction and interpretation (Figures 4A, S6, 3).

Model interpretation is enhanced by comparing the resulting cluster PSSMs with kinase specificity data such as PSPL to identify putative upstream kinases modulating signaling clusters. Computational validations showed that DDMC was able to correctly associate AKT1 and ERK2 with clusters of their respective substrates (Figure 3). It is worth noting, however, that kinase specificity is defined by additional features beyond the phosphosite motif such as kinase-substrate co-localization, regulation by phosphosite-binding domains (e.g., SH2, PTB domains), or docking. In addition, a major limitation of PSPL experiments is that since they do not provide docking information, the real affinity between the string of identified peptide residues as key determinants of specificity of a sequence motif and the interacting kinase domain is unknown. This limitation could also compromise kinase-cluster associations established by DDMC. A method combining bacterial surface-display of peptide libraries with next-generation sequencing tackles this limitation by quantifying the specificity of a kinase to virtually all possible motif combinations (*51*). Thus, as the number of profiled kinases with this technique increases, these measurements could be used to rank cluster peptides by magnitude of specificity to a specific kinase to make better upstream kinase predictions.

A key benefit of DDMC is that the identified clusters are not limited to pre-existing motifs and are therefore not dependent on prior experimentally validated kinase-substrate interactions. Thereby, this method could improve our understanding of the signaling effects of understudied kinases. For instance, our model predicts NEK1&2 promote, at least in part, a cluster with strikingly increased signaling in NATs compared to tumors. Further exploration of this cluster led us to hypothesize that the lack of NEK signaling in tumor samples might associated with the absence of ciliagenesis and adaptation to hypoxia in lung tumors (Figure 5G-H). Additionally, we show that cluster 8, which greatly contributes to explain the signaling differences between STK11 WT and mutant samples in tumors (Figure 6C), is enriched with proteins such as GOLPH3 and Myosin 18A that have been shown to promote Golgi fragmentation in cancer (*38–40*). This prompts us to consider the novel interaction between CK1 and these signaling molecules.

An additional major challenge being faced during the analysis of large-scale signaling data is missingness. Given that statistical tools often require complete data sets, researchers use standard methods to impute missing values such as the peptides’ mean or minimum signal, constant zero, or PCA imputation only in peptides wherein at least 50% of their samples were required to have non-missing values as excessive missing values can result in poor imputation (*10, 11, 52*). In this study we show that DDMC can model a data set of 30,561 peptides after filtering out any phosphosites that were not captured in at least 2 TMT (up to ~80% of missingness) by ignoring unobserved values during EM distribution estimation and calculation of GMM probabilities (see methods). Therefore, this method enables clustering of signaling data despite a remarkable number of missing values. Furthermore, DDMC clearly outperforms the imputation performance of using the peptides’ mean, minimum signal, or constant zero and provides similar results to PCA imputation. This important feature could offer the possibility of conducting pan-cancer phosphoproteomics studies using readily available large-scale clinical phosphoproteomic data.

The benefit of building algorithms combining different information sources is evident in previously published approaches. For instance, INKA predicts active kinases by integrating scores reflecting both phosphorylation status and substrate abundance (*53*). In another study, Exarchos et al. formulated a decision support system that integrates clinical, imaging, and genomic data to identify the factors that contribute to oral cancer progression and predict relapses. The authors found that combining the more accurate individual predictors yielded better predictions than those generated by other strategies reported in the literature (*54*). Finally, BOADICEA is a method that allows systematic risk stratification of breast cancer patients by incorporating the effects of lifestyle, hormonal and reproductive risk factors, mammographic density, and of the common breast cancer susceptibility genetic variants into the prediction model (55).

In total, in this study we show that combining the information about the sequence features and phosphorylation abundance leads to more robust clustering of global signaling measurements. Use of the DDMC clusters to regress against cell phenotypes led to enhanced model predictions and interpretation. Thus, we propose DDMC as a general and flexible strategy for phosphoproteomic analysis.

## Materials and Methods

All analysis was implemented in Python v3.9 and can be found at https://github.com/meyer-lab/resistance-MS.

### Expectation-maximization (EM) algorithm architecture

We constructed a modified mixture model that clusters peptides based on both their abundance across conditions and sequence. The model is defined by a given number of clusters and weighting factor to prioritize either the data or the sequence information. Fitting was performed using expectation-maximization, initialized at a starting point. The starting point was derived from k-means clustering the abundance data after missing values were imputed by PCA with a component number equal to the number of clusters. During the expectation (E) step, the algorithm calculates the probability of each peptide being assigned to each cluster. In the maximization (M) step, each cluster’s distributions are fit using the weighted cluster assignments. The peptide sequence and abundance assignments within the E step are combined by taking the sum of the log-likelihood of both assignments. The peptide log-likelihood is multiplied by the user-defined weighting factor immediately before to influence its importance. Both steps repeat until convergence as defined by the increase in model log-likelihood between iterations falling below a user-defined threshold.

### Phosphorylation site abundance clustering in the presence of missing values

We modeled the log-transformed abundance of each phosphopeptide as following a multivariate Gaussian distribution with diagonal covariance. Each dimension of this distribution represents the abundance of that peptide within a given sample. For example, within a data set of 100 patients and 1000 peptides, using 10 clusters, the data is represented by 10 Gaussian distributions of 100 dimensions. Unobserved/missing values were indicated as NaN and ignored during both distribution estimation and when calculating probabilities. Any peptides that were detected in only one TMT experiment were discarded.

### Sequence-cluster comparison

#### PAM250

During model initialization, the pairwise distance between all peptides in the dataset was calculated using the PAM250 matrix. The mean distance from each peptide to a given cluster could then be calculated by:

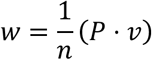

Where *P* is the *n* × *n* distance matrix, *n* is the number of peptides in the dataset, *v* is the probability of each peptide being assigned to the cluster of interest, and *w* is the log-probabilities of cluster assignment.

#### Binomial enrichment

We alternatively used a binomial enrichment model for the sequence representation of a cluster based on earlier work (*55*). Upon model initialization, a background matrix *i* × *j* × *k* was created with a position-specific scoring matrix of all the sequences together. Next, an *T* data tensor *i* was created where *j* is the number of peptides, *k* is the number of amino acid possibilities, and *k* is the position relative to the phosphorylation site. This tensor contained 1 where an amino acid was present for that position and peptide, and 0 elsewhere.

Within each iteration, the cluster motif would be updated using *v*, the probability of each peptide being assigned to the cluster of interest. First, a weighted count for each amino acid and position would be assembled:

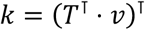

Because peptides can be partially assigned to a cluster, the counts of each amino acid and position can take continuous values. We therefore generalized the binomial distribution to allow continuous values using the regularized incomplete Beta function:

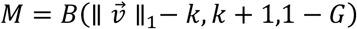

Finally, the log-probability of membership for each peptide was calculated based on the product of each amino acid-position probability.

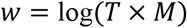

We confirmed that this provided identical results to a binomial enrichment model for integer counts of amino acids (*55*) but allowed for partial assignment of peptides to clusters.

### Quantifying the influence of sequence versus data

The magnitude of the weight used to scale the sequence and data scores is arbitrary. We do know that with a weight of 0 the model only uses the phosphorylation measurements. Alternatively, with an enormously large weight the motif information is prioritized. However, we do not know to what extent each information source is prioritized in general. Therefore, to quantify the relative importance of each type of data, we calculated our clustering results at each weighting extreme, and then calculated the Frobenius norm of the resulting peptide assignments between those and the clustering of interest.

### Generating Cluster Motifs and Upstream Kinase Predictions

For each cluster we computed a position-specific-scoring matrix (PSSM). To do so, we populated a residue/position matrix with the sum of the corresponding cluster probabilities for every peptide. Once all peptides were accounted for, the resulting matrix was normalized by averaging the mean probability across amino acids and log2-transformed to generate a PSSM. In parallel, we computed a PSSM including all sequences that served as background to account for the different amino acid occurrences within the data set. Then, we subtracted each cluster PSSM with the background PSSM and limited any large negative numbers to −3. Next, we extracted several kinase specificity profiling results from the literature (*16, 18, 18, 19*). The distance between PSSM and PSSL motifs was calculated using by the Frobenius norm of the difference. Motif logo plots were generated using logomaker (*56*).

### Evaluate clustering by imputation of values

To evaluate the ability of our model to handle missing values, we removed random, individual TMT experiments for each peptide and used the model to impute these values. The number of missing values per peptide is highly variable. Therefore, in our error quantitation, we stratified peptides by their missingness percentage and computed the average mean squared error between the actual values and predictions—or imputed peptide average—in each group. We calculated the reconstruction error across different combinations of cluster numbers and weights using the same process.

### Associating clusters with molecular and clinical features

To find clusters that tracked with specific molecular or clinical features we implemented two different strategies: logistic regression and hypothesis testing. For binary problems such as Tumor vs NAT samples or mutational status we used l1-regularized logistic regression and Mann-Whitney rank tests. In the former, we tried to predict the feature of interest using the phosphorylation signal of the cluster centers, whereas in the latter, for each cluster we split all patients according to their specific feature and tested whether the difference in the median signal between both groups was statistically different. We performed Bonferroni correction on the p-values computed by the Mann-Whitney tests. Gene ontology analysis was performed using the GENEONTOLOGY software (geneontology.org) (*57, 58*).

## Author contributions

A.S.M. conceived the project. Both authors performed the analysis. Both authors wrote the manuscript.

## References

1. T. Hunter, Protein kinases and phosphatases: the yin and yang of protein phosphorylation and signaling. Cell. 80, 225–36 (1995).

2. M. B. Yaffe, Why geneticists stole cancer research even though cancer is primarily a signaling disease. Sci Signal. 12 (2019), doi:10.1126/scisignal.aaw3483.

3. P. Casado, J.-C. Rodriguez-Prados, S. C. Cosulich, S. Guichard, B. Vanhaesebroeck, S. Joel, P. R. Cutillas, Kinase-substrate enrichment analysis provides insights into the heterogeneity of signaling pathway activation in leukemia cells. Sci Signal. 6, rs6 (2013).

4. R. Beekhof, C. van Alphen, A. A. Henneman, J. C. Knol, T. V. Pham, F. Rolfs, M. Labots, E. Henneberry, T. Y. Le Large, R. R. de Haas, S. R. Piersma, V. Vurchio, A. Bertotti, L. Trusolino, H. M. Verheul, C. R. Jimenez, INKA, an integrative data analysis pipeline for phosphoproteomic inference of active kinases. Mol Syst Biol. 15, e8981 (2019).

5. P. V. Hornbeck, J. M. Kornhauser, V. Latham, B. Murray, V. Nandhikonda, A. Nord, E. Skrzypek, T. Wheeler, B. Zhang, F. Gnad, 15 years of PhosphoSitePlus®: integrating post-translationally modified sites, disease variants and isoforms. Nucleic Acids Res. 47, D433–D441 (2019).

6. R. Linding, L. J. Jensen, G. J. Ostheimer, M. A. T. M. van Vugt, C. Jørgensen, I. M. Miron, F. Diella, K. Colwill, L. Taylor, K. Elder, P. Metalnikov, V. Nguyen, A. Pasculescu, J. Jin, J. G. Park, L. D. Samson, J. R. Woodgett, R. B. Russell, P. Bork, M. B. Yaffe, T. Pawson, Systematic discovery of in vivo phosphorylation networks. Cell. 129, 1415–26 (2007).

7. J. C. Obenauer, L. C. Cantley, M. B. Yaffe, Scansite 2.0: Proteome-wide prediction of cell signaling interactions using short sequence motifs. Nucleic Acids Res. 31, 3635–41 (2003).

8. E. J. Needham, B. L. Parker, T. Burykin, D. E. James, S. J. Humphrey, Illuminating the dark phosphoproteome. Sci Signal. 12 (2019), doi:10.1126/scisignal.aau8645.

9. D. L. Tabb, L. Vega-Montoto, P. A. Rudnick, A. M. Variyath, A.-J. L. Ham, D. M. Bunk, L. E. Kilpatrick, D. D. Billheimer, R. K. Blackman, H. L. Cardasis, S. A. Carr, K. R. Clauser, J. D. Jaffe, K. A. Kowalski, T. A. Neubert, F. E. Regnier, B. Schilling, T. J. Tegeler, M. Wang, P. Wang, J. R. Whiteaker, L. J. Zimmerman, S. J. Fisher, B. W. Gibson, C. R. Kinsinger, M. Mesri, H. Rodriguez, S. E. Stein, P. Tempst, A. G. Paulovich, D. C. Liebler, C. Spiegelman, Repeatability and reproducibility in proteomic identifications by liquid chromatography-tandem mass spectrometry. J Proteome Res. 9, 761–76 (2010).

10. Y.-J. Chen, T. I. Roumeliotis, Y.-H. Chang, C.-T. Chen, C.-L. Han, M.-H. Lin, H.-W. Chen, G.-C. Chang, Y.-L. Chang, C.-T. Wu, M.-W. Lin, M.-S. Hsieh, Y.-T. Wang, Y.-R. Chen, I. Jonassen, F. Z. Ghavidel, Z.-S. Lin, K.-T. Lin, C.-W. Chen, P.-Y. Sheu, C.-T. Hung, K.-C. Huang, H.-C. Yang, P.-Y. Lin, T.-C. Yen, Y.-W. Lin, J.-H. Wang, L. Raghav, C.-Y. Lin, Y.-S. Chen, P.-S. Wu, C.-T. Lai, S.-H. Weng, K.-Y. Su, W.-H. Chang, P.-Y. Tsai, A. I. Robles, H. Rodriguez, Y.-J. Hsiao, W.-H. Chang, T.-Y. Sung, J.-S. Chen, S.-L. Yu, J. S. Choudhary, H.-Y. Chen, P.-C. Yang, Y.-J. Chen, Proteogenomics of Non-smoking Lung Cancer in East Asia Delineates Molecular Signatures of Pathogenesis and Progression. Cell. 182, 226–244.e17 (2020).

11. M. A. Gillette, S. Satpathy, S. Cao, S. M. Dhanasekaran, S. V. Vasaikar, K. Krug, F. Petralia, Y. Li, W.-W. Liang, B. Reva, A. Krek, J. Ji, X. Song, W. Liu, R. Hong, L. Yao, L. Blumenberg, S. R. Savage, M. C. Wendl, B. Wen, K. Li, L. C. Tang, M. A. MacMullan, S. C. Avanessian, M. H. Kane, C. J. Newton, M. Cornwell, R. B. Kothadia, W. Ma, S. Yoo, R. Mannan, P. Vats, C. Kumar-Sinha, E. A. Kawaler, T. Omelchenko, A. Colaprico, Y. Geffen, Y. E. Maruvka, F. da Veiga Leprevost, M. Wiznerowicz, Z. H. Gümüş, R. R. Veluswamy, G. Hostetter, D. I. Heiman, M. A. Wyczalkowski, T. Hiltke, M. Mesri, C. R. Kinsinger, E. S. Boja, G. S. Omenn, A. M. Chinnaiyan, H. Rodriguez, Q. K. Li, S. D. Jewell, M. Thiagarajan, G. Getz, B. Zhang, D. Fenyö, K. V. Ruggles, M. P. Cieslik, A. I. Robles, K. R. Clauser, R. Govindan, P. Wang, A. I. Nesvizhskii, L. Ding, D. R. Mani, S. A. Carr,, Proteogenomic Characterization Reveals Therapeutic Vulnerabilities in Lung Adenocarcinoma. Cell. 182, 200–225.e35 (2020).

12. A. Zarrinpar, S.-H. Park, W. A. Lim, Optimization of specificity in a cellular protein interaction network by negative selection. Nature. 426, 676–80 (2003).

13. C. S. H. Tan, A. Pasculescu, W. A. Lim, T. Pawson, G. D. Bader, R. Linding, Positive selection of tyrosine loss in metazoan evolution. Science. 325, 1686–8 (2009).

14. T. Obata, M. B. Yaffe, G. G. Leparc, E. T. Piro, H. Maegawa, A. Kashiwagi, R. Kikkawa, L. C. Cantley, Peptide and protein library screening defines optimal substrate motifs for AKT/PKB. J Biol Chem. 275, 36108–15 (2000).

15. J. Mok, P. M. Kim, H. Y. K. Lam, S. Piccirillo, X. Zhou, G. R. Jeschke, D. L. Sheridan, S. A. Parker, V. Desai, M. Jwa, E. Cameroni, H. Niu, M. Good, A. Remenyi, J.-L. N. Ma, Y.-J. Sheu, H. E. Sassi, R. Sopko, C. S. M. Chan, C. De Virgilio, N. M. Hollingsworth, W. A. Lim, D. F. Stern, B. Stillman, B. J. Andrews, M. B. Gerstein, M. Snyder, B. E. Turk, Deciphering protein kinase specificity through large-scale analysis of yeast phosphorylation site motifs. Sci Signal. 3, ra12 (2010).

16. B. van de Kooij, P. Creixell, A. van Vlimmeren, B. A. Joughin, C. J. Miller, N. Haider, C. D. Simpson, R. Linding, V. Stambolic, B. E. Turk, M. B. Yaffe, Comprehensive substrate specificity profiling of the human Nek kinome reveals unexpected signaling outputs. Elife. 8 (2019), doi:10.7554/elife.44635.

17. J. E. Hutti, E. T. Jarrell, J. D. Chang, D. W. Abbott, P. Storz, A. Toker, L. C. Cantley, B. E. Turk, A rapid method for determining protein kinase phosphorylation specificity. Nat Methods. 1, 27–9 (2004).

18. M. J. Begley, C.-h. Yun, C. A. Gewinner, J. M. Asara, J. L. Johnson, A. J. Coyle, M. J. Eck, I. Apostolou, L. C. Cantley, EGF-receptor specificity for phosphotyrosine-primed substrates provides signal integration with Src. Nat Struct Mol Biol. 22, 983–90 (2015).

19. M. L. Miller, L. J. Jensen, F. Diella, C. Jørgensen, M. Tinti, L. Li, M. Hsiung, S. A. Parker, J. Bordeaux, T. Sicheritz-Ponten, M. Olhovsky, A. Pasculescu, J. Alexander, S. Knapp, N. Blom, P. Bork, S. Li, G. Cesareni, T. Pawson, B. E. Turk, M. B. Yaffe, S. Brunak, R. Linding, Linear motif atlas for phosphorylation-dependent signaling. Sci Signal. 1, ra2 (2008).

20. M. Hijazi, R. Smith, V. Rajeeve, C. Bessant, P. R. Cutillas, Reconstructing kinase network topologies from phosphoproteomics data reveals cancer-associated rewiring. Nat Biotechnol. 38, 493–502 (2020).

21. S. M. Carlson, C. R. Chouinard, A. Labadorf, C. J. Lam, K. Schmelzle, E. Fraenkel, F. M. White, Large-scale discovery of ERK2 substrates identifies ERK-mediated transcriptional regulation by ETV3. Sci Signal. 4, rs11 (2011).

22. L. Moniz, P. Dutt, N. Haider, V. Stambolic, Nek family of kinases in cell cycle, checkpoint control and cancer. Cell Div. 6, 18 (2011).

23. G. V. Meirelles, A. M. Perez, E. E. de Souza, F. L. Basei, P. F. Papa, T. D. Melo Hanchuk, V. B. Cardoso, J. Kobarg, “Stop Ne(c)king around”: How interactomics contributes to functionally characterize Nek family kinases. World J Biol Chem. 5, 141–60 (2014).

24. L. Fabbri, F. Bost, N. M. Mazure, Primary Cilium in Cancer Hallmarks. Int J Mol Sci. 20 (2019), doi:10.3390/ijms20061336.

25. O. V. Plotnikova, E. A. Golemis, E. N. Pugacheva, Cell cycle-dependent ciliogenesis and cancer. Cancer Res. 68, 2058–61 (2008).

26. R. M. Brosh, DNA helicases involved in DNA repair and their roles in cancer. Nat Rev Cancer. 13, 542–58 (2013).

27. J. So, A. Pasculescu, A. Y. Dai, K. Williton, A. James, V. Nguyen, P. Creixell, E. M. Schoof, J. Sinclair, M. Barrios-Rodiles, J. Gu, A. Krizus, R. Williams, M. Olhovsky, J. W. Dennis, J. L. Wrana, R. Linding, C. Jorgensen, T. Pawson, K. Colwill, Integrative analysis of kinase networks in TRAIL-induced apoptosis provides a source of potential targets for combination therapy. Sci Signal. 8, rs3 (2015).

28. M. V. Bennetzen, D. H. Larsen, J. Bunkenborg, J. Bartek, J. Lukas, J. S. Andersen, Site-specific phosphorylation dynamics of the nuclear proteome during the DNA damage response. Mol Cell Proteomics. 9, 1314–23 (2010).

29. A. J. Rabalski, L. Gyenis, D. W. Litchfield, Molecular Pathways: Emergence of Protein Kinase CK2 (CSNK2) as a Potential Target to Inhibit Survival and DNA Damage Response and Repair Pathways in Cancer Cells. Clin Cancer Res. 22, 2840–7 (2016).

30. A. Siddiqui-Jain, J. Bliesath, D. Macalino, M. Omori, N. Huser, N. Streiner, C. B. Ho, K. Anderes, C. Proffitt, S. E. O’Brien, J. K. C. Lim, D. D. Von Hoff, D. M. Ryckman, W. G. Rice, D. Drygin, CK2 inhibitor CX-4945 suppresses DNA repair response triggered by DNA-targeted anticancer drugs and augments efficacy: mechanistic rationale for drug combination therapy. Mol Cancer Ther. 11, 994–1005 (2012).

31. H. Ji, M. R. Ramsey, D. N. Hayes, C. Fan, K. McNamara, P. Kozlowski, C. Torrice, M. C. Wu, T. Shimamura, S. A. Perera, M.-C. Liang, D. Cai, G. N. Naumov, L. Bao, C. M. Contreras, D. Li, L. Chen, J. Krishnamurthy, J. Koivunen, L. R. Chirieac, R. F. Padera, R. T. Bronson, N. I. Lindeman, D. C. Christiani, X. Lin, G. I. Shapiro, P. A. Jänne, B. E. Johnson, M. Meyerson, D. J. Kwiatkowski, D. H. Castrillon, N. Bardeesy, N. E. Sharpless, K.-K. Wong, LKB1 modulates lung cancer differentiation and metastasis. Nature. 448, 807–10 (2007).

32. A. L. Manning, M. S. Longworth, N. J. Dyson, Loss of pRB causes centromere dysfunction and chromosomal instability. Genes Dev. 24, 1364–76 (2010).

33. A. L. Manning, S. A. Yazinski, B. Nicolay, A. Bryll, L. Zou, N. J. Dyson, Suppression of genome instability in pRB-deficient cells by enhancement of chromosome cohesion. Mol Cell. 53, 993–1004 (2014).

34. M.-H. Nguyen, J. Koinuma, K. Ueda, T. Ito, E. Tsuchiya, Y. Nakamura, Y. Daigo, Phosphorylation and activation of cell division cycle associated 5 by mitogen-activated protein kinase play a crucial role in human lung carcinogenesis. Cancer Res. 70, 5337–47 (2010).

35. R. H. Hruban, M. I. Canto, M. Goggins, R. Schulick, A. P. Klein, Update on familial pancreatic cancer. Adv Surg. 44, 293–311 (2010).

36. R. K. Gill, S.-H. Yang, D. Meerzaman, L. E. Mechanic, E. D. Bowman, H.-S. Jeon, S. Roy Chowdhuri, A. Shakoori, T. Dracheva, K.-M. Hong, J. Fukuoka, J.-H. Zhang, C. C. Harris, J. Jen, Frequent homozygous deletion of the LKB1/STK11 gene in non-small cell lung cancer. Oncogene. 30, 3784–91 (2011).

37. A. Petrosyan, Onco-Golgi: Is Fragmentation a Gate to Cancer Progression? Biochem Mol Biol J. 1 (2015), doi:10.21767/2471-8084.100006.

38. X. Hua, L. Yu, W. Pan, X. Huang, Z. Liao, Q. Xian, L. Fang, H. Shen, Increased expression of Golgi phosphoprotein-3 is associated with tumor aggressiveness and poor prognosis of prostate cancer. Diagn Pathol. 7, 127 (2012).

39. Z. Zeng, H. Lin, X. Zhao, G. Liu, X. Wang, R. Xu, K. Chen, J. Li, L. Song, Overexpression of GOLPH3 promotes proliferation and tumorigenicity in breast cancer via suppression of the FOXO1 transcription factor. Clin Cancer Res. 18, 4059–69 (2012).

40. B.-S. Hu, H. Hu, C.-Y. Zhu, Y.-L. Gu, J.-P. Li, Overexpression of GOLPH3 is associated with poor clinical outcome in gastric cancer. Tumour Biol. 34, 515–20 (2012).

41. T. Sasaki, J. Koivunen, A. Ogino, M. Yanagita, S. Nikiforow, W. Zheng, C. Lathan, J. P. Marcoux, J. Du, K. Okuda, M. Capelletti, T. Shimamura, D. Ercan, M. Stumpfova, Y. Xiao, S. Weremowicz, M. Butaney, S. Heon, K. Wilner, J. G. Christensen, M. J. Eck, K.-K. Wong, N. Lindeman, N. S. Gray, S. J. Rodig, P. A. Jänne, A novel ALK secondary mutation and EGFR signaling cause resistance to ALK kinase inhibitors. Cancer Res. 71, 6051–60 (2011).

42. M. Miyawaki, H. Yasuda, T. Tani, J. Hamamoto, D. Arai, K. Ishioka, K. Ohgino, S. Nukaga, T. Hirano, I. Kawada, K. Naoki, Y. Hayashi, T. Betsuyaku, K. Soejima, Overcoming EGFR Bypass Signal-Induced Acquired Resistance to ALK Tyrosine Kinase Inhibitors in ALK-Translocated Lung Cancer. Mol Cancer Res. 15, 106–114 (2016).

43. D. Aran, Z. Hu, A. J. Butte, xCell: digitally portraying the tissue cellular heterogeneity landscape. Genome Biol. 18, 220 (2017).

44. A. I. Khan, S. M. Kerfoot, B. Heit, L. Liu, G. Andonegui, B. Ruffell, P. Johnson, P. Kubes, Role of CD44 and hyaluronan in neutrophil recruitment. J Immunol. 173, 7594–601 (2004).

45. J. D. Klement, A. V. Paschall, P. S. Redd, M. L. Ibrahim, C. Lu, D. Yang, E. Celis, S. I. Abrams, K. Ozato, K. Liu, An osteopontin/CD44 immune checkpoint controls CD8+ T cell activation and tumor immune evasion. J Clin Invest. 128, 5549–5560 (2018).

46. C. M. Porter, M. A. Havens, N. A. Clipstone, Identification of amino acid residues and protein kinases involved in the regulation of NFATc subcellular localization. J Biol Chem. 275, 3543–51 (2000).

47. W. Yang, S. A. Gibson, Z. Yan, H. Wei, J. Tao, B. Sha, H. Qin, E. N. Benveniste, Protein kinase 2 (CK2) controls CD4. Mucosal Immunol. 13, 788–798 (2020).

48. S. Martínez-Høyer, A. Aranguren-Ibáñez, J. García-García, E. Serrano-Candelas, J. Vilardell, V. Nunes, F. Aguado, B. Oliva, E. Itarte, M. Pérez-Riba, Protein kinase CK2-dependent phosphorylation of the human Regulators of Calcineurin reveals a novel mechanism regulating the calcineurin-NFATc signaling pathway. Biochim Biophys Acta. 1833, 2311–21 (2013).

49. F. Skoulidis, M. E. Goldberg, D. M. Greenawalt, M. D. Hellmann, M. M. Awad, J. F. Gainor, A. B. Schrock, R. J. Hartmaier, S. E. Trabucco, L. Gay, S. M. Ali, J. A. Elvin, G. Singal, J. S. Ross, D. Fabrizio, P. M. Szabo, H. Chang, A. Sasson, S. Srinivasan, S. Kirov, J. Szustakowski, P. Vitazka, R. Edwards, J. A. Bufill, N. Sharma, S.-H. I. Ou, N. Peled, D. R. Spigel, H. Rizvi, E. J. Aguilar, B. W. Carter, J. Erasmus, D. F. Halpenny, A. J. Plodkowski, N. M. Long, M. Nishino, W. L. Denning, A. Galan-Cobo, H. Hamdi, T. Hirz, P. Tong, J. Wang, J. Rodriguez-Canales, P. A. Villalobos, E. R. Parra, N. Kalhor, L. M. Sholl, J. L. Sauter, A. A. Jungbluth, M. Mino-Kenudson, R. Azimi, Y. Y. Elamin, J. Zhang, G. C. Leonardi, F. Jiang, K.-K. Wong, J. J. Lee, V. A. Papadimitrakopoulou, I. I. Wistuba, V. A. Miller, G. M. Frampton, J. D. Wolchok, A. T. Shaw, P. A. Jänne, P. J. Stephens, C. M. Rudin, W. J. Geese, L. A. Albacker, J. V. Heymach, Cancer Discov. 8, 822–835 (2018).

50. K. Kourou, T. P. Exarchos, K. P. Exarchos, M. V. Karamouzis, D. I. Fotiadis, Machine learning applications in cancer prognosis and prediction. Comput Struct Biotechnol J. 13, 8–17 (2014).

51. N. H. Shah, M. Löbel, A. Weiss, J. Kuriyan, Fine-tuning of substrate preferences of the Src-family kinase Lck revealed through a high-throughput specificity screen. Elife. 7 (2018), doi:10.7554/elife.35190.

52. B. Deb, P. Sengupta, J. Sambath, P. Kumar, Bioinformatics Analysis of Global Proteomic and Phosphoproteomic Data Sets Revealed Activation of NEK2 and AURKA in Cancers. Biomolecules. 10 (2020), doi:10.3390/biom10020237.

53. R. Beekhof, C. van Alphen, A. A. Henneman, J. C. Knol, T. V. Pham, F. Rolfs, M. Labots, E. Henneberry, T. Y. Le Large, R. R. de Haas, S. R. Piersma, V. Vurchio, A. Bertotti, L. Trusolino, H. M. Verheul, C. R. Jimenez, INKA, an integrative data analysis pipeline for phosphoproteomic inference of active kinases. Mol Syst Biol. 15, e8250 (2019).

54. K. P. Exarchos, Y. Goletsis, D. I. Fotiadis, Multiparametric decision support system for the prediction of oral cancer reoccurrence. IEEE Trans Inf Technol Biomed. 16, 1127–34 (2011).

55. A. Antoniou, A. Cunningham, J. Peto, D. G Evans, F Lalloo, S. A. Narod, H. A. Risch, J.E. Eyfjord, J.L. Hopper, M. C. Southey, H. Olsson, O. Johannsson, A. Borg, B. Passini, P. Radice, S Manoukian, D. M Eccles, N. Tang, E. Olah, H. Anton-Culver, E. Warner, J. Lubinski, J. Gronwald, B. Gorski, L. Tryggvadottir, K. Syrjakoski, O.P. Kallioniemi, H. Eerola, H. Nevanlinna, P.D.P Pharoah, D.F. Easton, The BOADICEA model of genetic susceptibility to breast and ovarian cancers: Updates and extensions. British Journal of Cancer. 8, 1457–1466 (2008).

55. D. Schwartz, S. P. Gygi, An iterative statistical approach to the identification of protein phosphorylation motifs from large-scale data sets. Nat Biotechnol. 23, 1391–8 (2005).

56. A. Tareen, J. B. Kinney, Logomaker: Beautiful sequence logos in python. Cold Spring Harbor Laboratory (2019), doi:10.1101/635029.

57. M. Ashburner, C. A. Ball, J. A. Blake, D. Botstein, H. Butler, J. M. Cherry, A. P. Davis, K. Dolinski, S. S. Dwight, J. T. Eppig, M. A. Harris, D. P. Hill, L. Issel-Tarver, A. Kasarskis, S. Lewis, J. C. Matese, J. E. Richardson, M. Ringwald, G. M. Rubin, G. Sherlock, Gene ontology: tool for the unification of biology. The Gene Ontology Consortium. Nat Genet. 25, 25–9 (2000).

58. The Gene Ontology resource: enriching a GOld mine. Nucleic Acids Res. 49, D325–D334 (2021).

