## Supplement for "Motif-based phosphoproteome clustering improves modeling and interpretation"

### Supplementary Materials

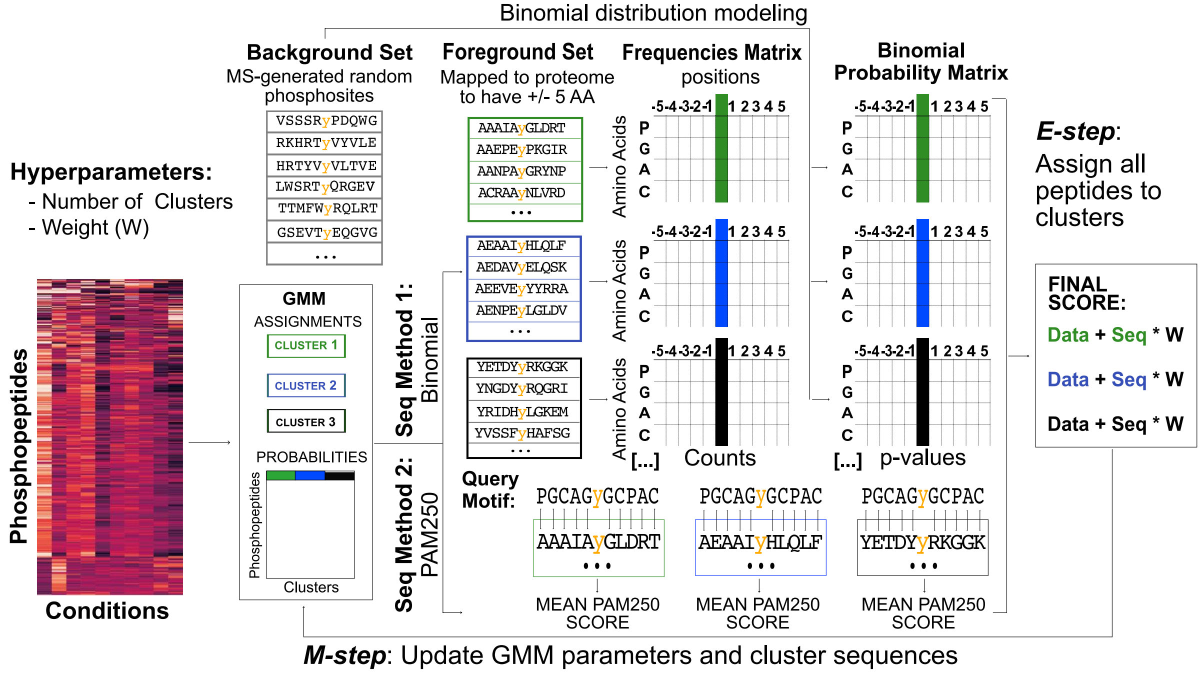


**Figure S1: Schematic of the DDMC simultaneous data and peptide sequence clustering approach.** Peptides are initialized into clusters at random. The measurements of abundance are represented by a multivariate Gaussian mixture model where each dimension of the distribution represents the abundance within a sample. Next, an expectation-maximization fitting scheme is used. During the expectation step, the distance of each peptide sequence to each cluster is calculated. This is done either through a binomial enrichment scheme (method 1) or using the average PAM250 distance (method 2). In parallel, the distance of each peptide abundance is compared to the cluster centers. These two distances are combined to update the assignments of each peptide to each cluster. During the maximization step, the cluster centers of the data are updated based on the weighted average of the peptide abundances in each condition. The peptide motifs are similarly updated through a weighted combination of the assigned peptides. Both steps continue sequentially until the change in peptide assignments between each iteration drops below a threshold.

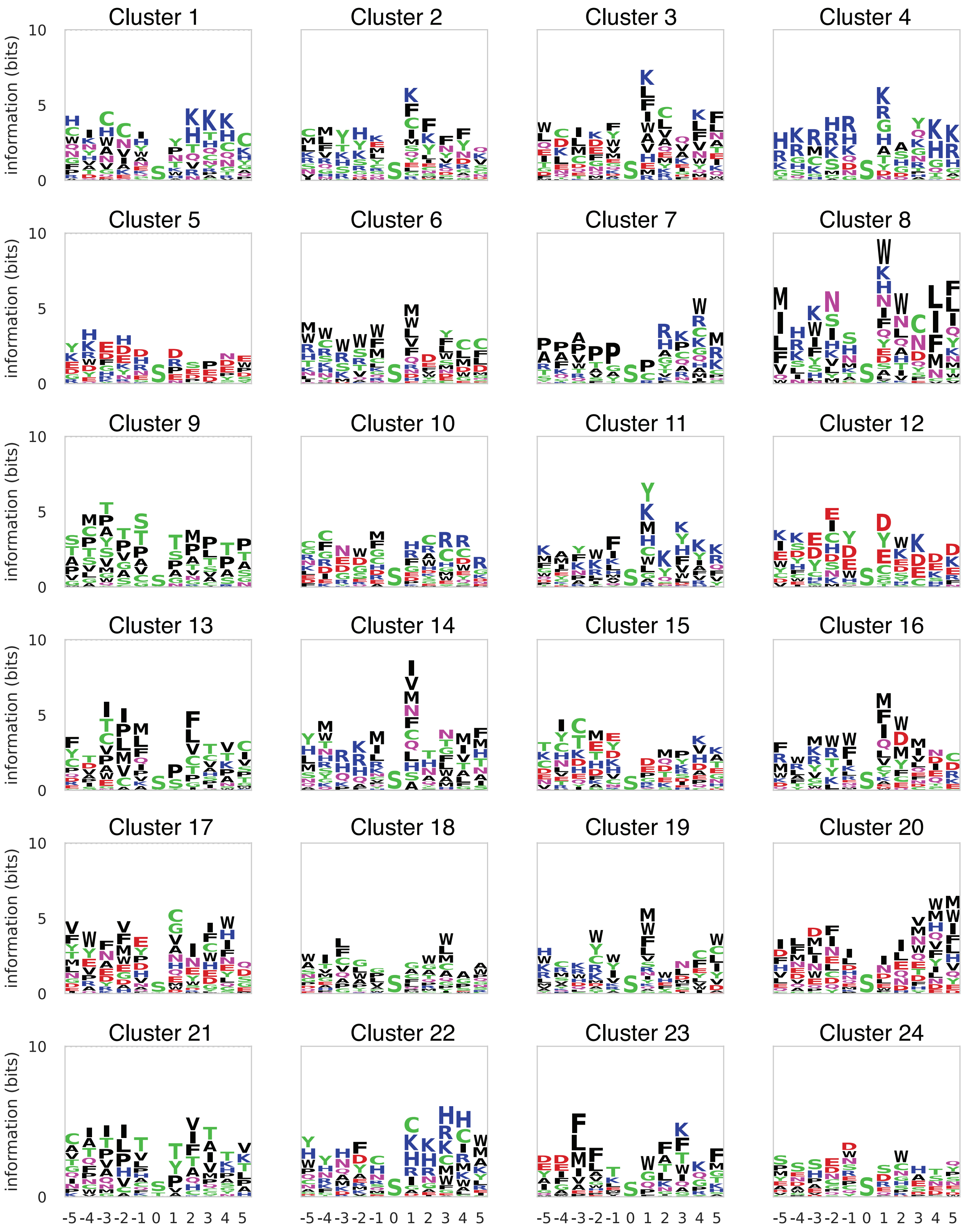


**Figure S2: Logo plots of all Cluster PSSMs.** (A-U) Sequence motifs of clusters 1 through 24.

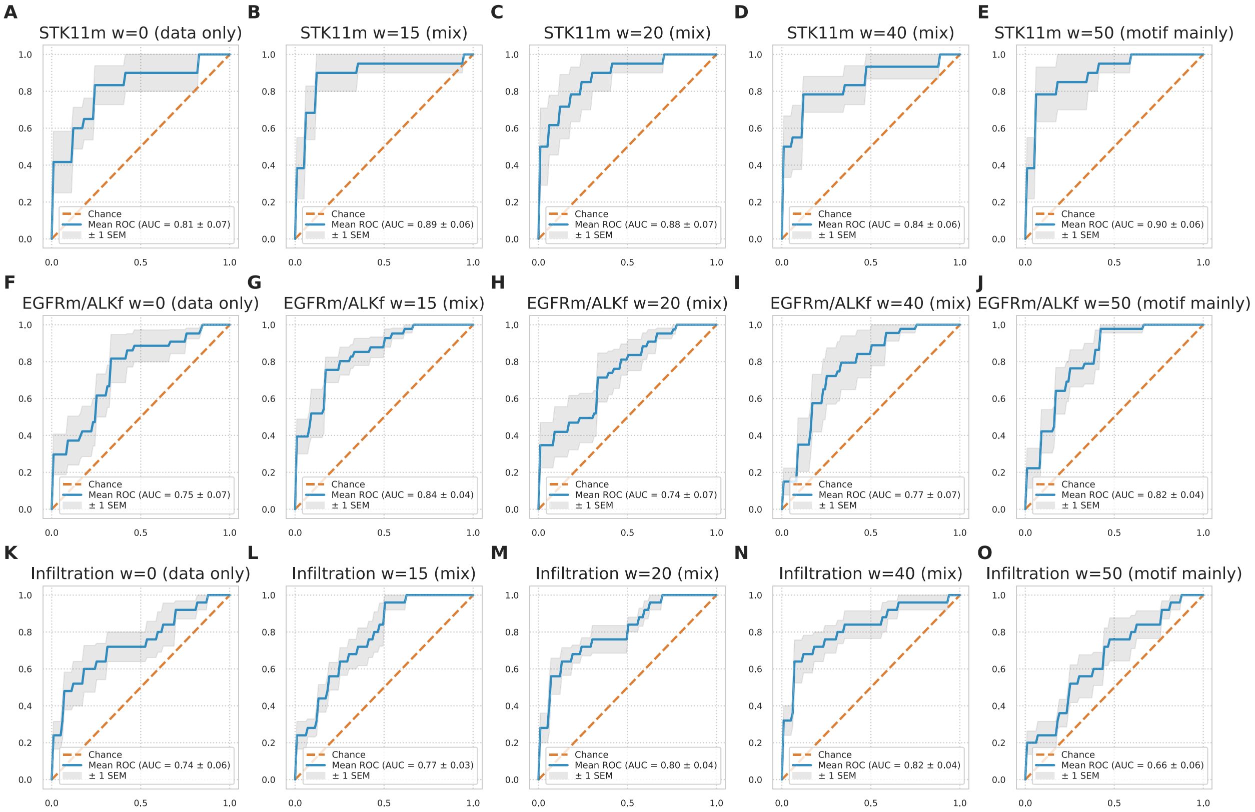


**Figure S3: Sequence information enhances model prediction.** (A-O) Performance of a regression model predicting the mutational status of STK11 (A-E) EGFR and/or ALK (F-J) and tumor infiltration level (hot versus cold) (K-O) in LUAD patients using either only phosphorylation data (0), mainly peptide sequences (50), or a mix (15, 20, 40). EGFRm/ALKf; EGFR mutant and/or ALK fusion.

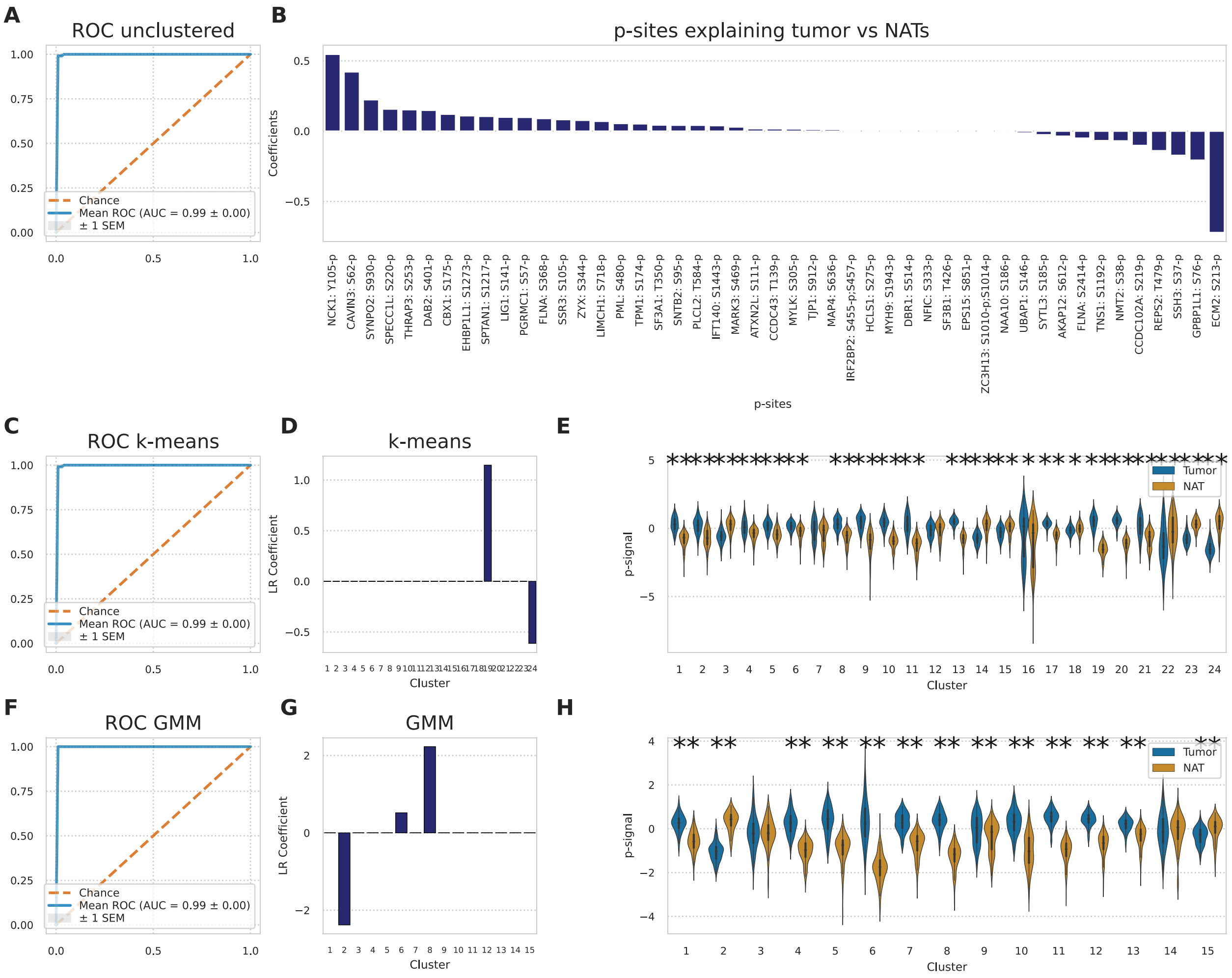


**Figure S4: Additional modeling strategies to find conserved tumor differences compared to NATs.** A) ROC plot of a logistic regression model fit to the complete portion of the phosphoproteomic CPTAC LUAD data set. B) Phospho-peptides with largest weights explaining the observed differences between tumors and NATs. C-D) ROC plot of a logistic regression model fit to the complete portion of the signaling data set clustered by k-means (D) and its corresponding cluster coefficients (D). E) p-site abundance between NAT and tumor patients per k-means cluster and its statistical significance. (F-H) Analysis using GMM clustering. (* = p-value < 0.05 and ** = p-value < 0.001)

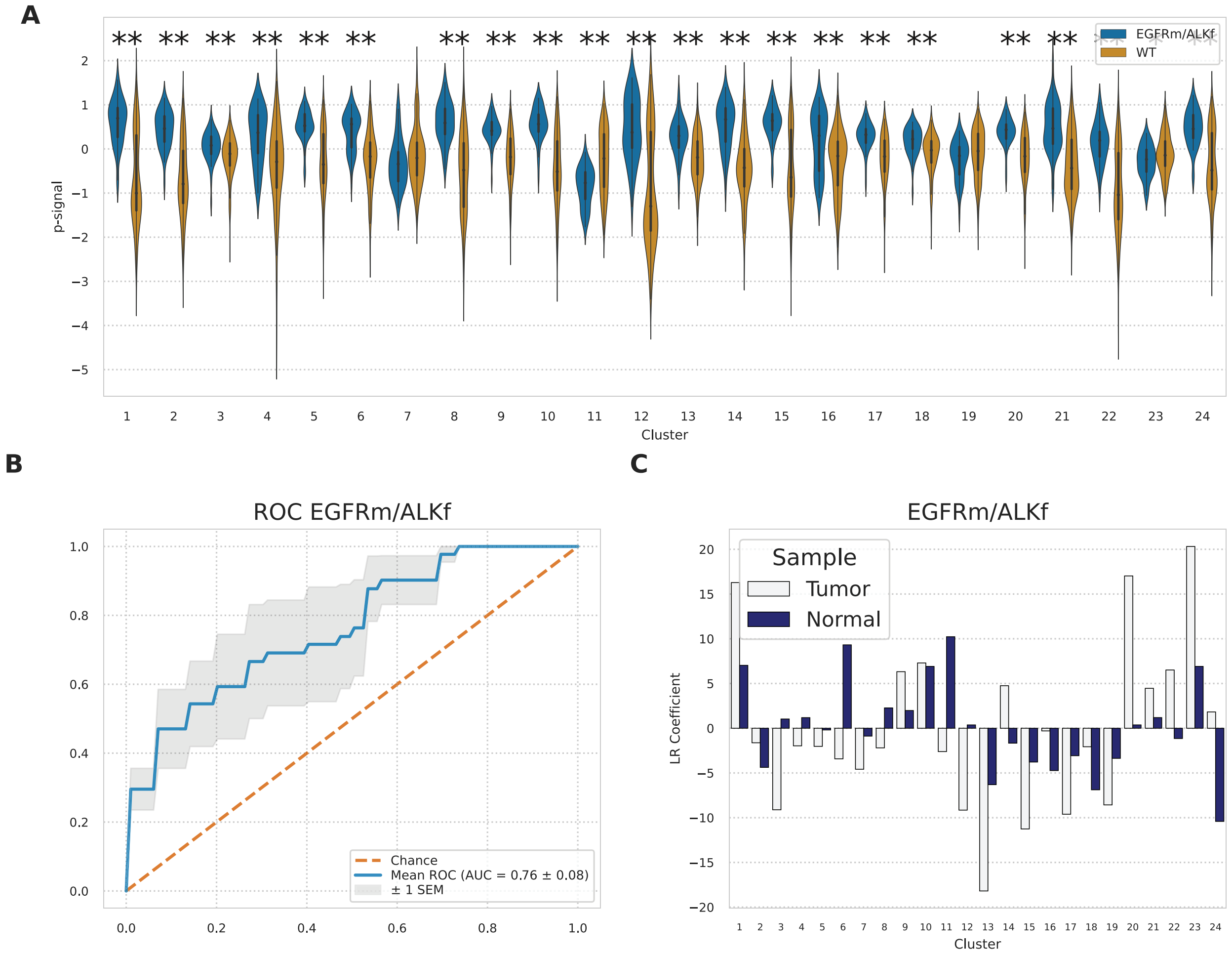


**Figure S5: Prediction of patient samples harboring EGFR mutations or ALK fusions (EGFRm/ALKf).** A) Phosphorylation signal of DDMC clusters grouped by EGFRm/ALKf mutant and WT samples. Its statistical significance is indicated on the top part of the plot via a series of Mann Whitney rank test. (* = p-value < 0.05 and ** = p-value < 0.001) B) ROC plot of the logistic regression model fit to the DDMC clusters. (D-G) Logo plots of the sequence motifs for clusters 2, 13, 20, and 23. H) Prediction of upstream kinases corresponding to the aforementioned clusters.

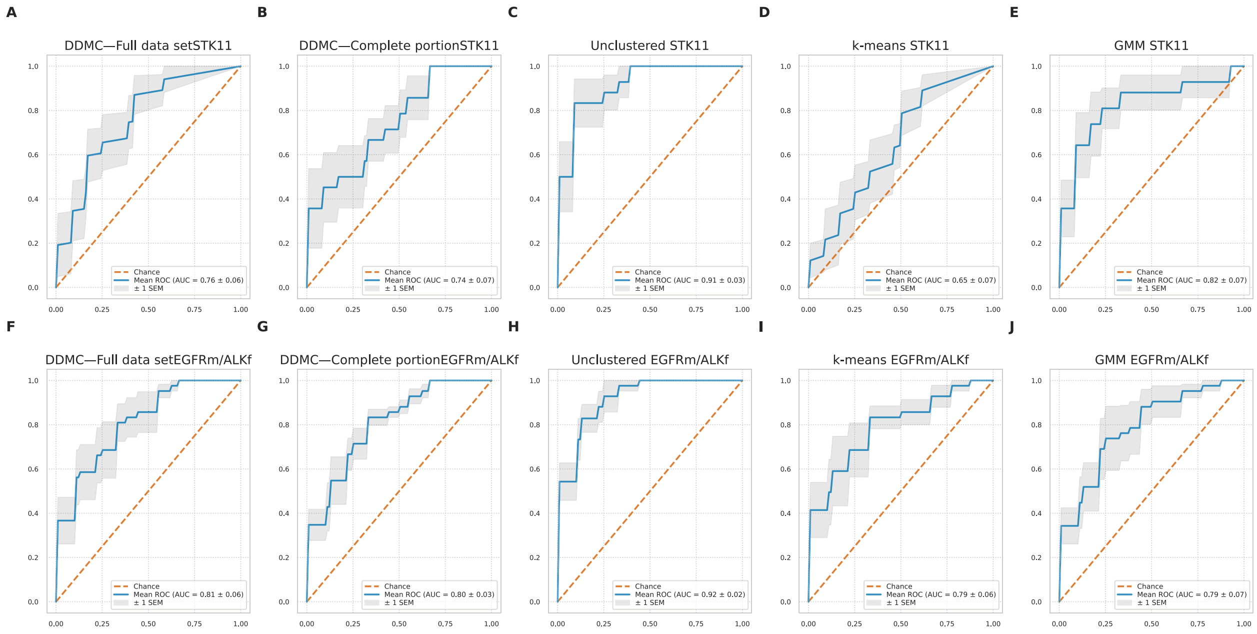


**Figure S6: Comparing the predictive performance of a logistic regression model using different modeling strategies.** (A-D) Performance of a logistic regression model predicting STK11 mutational status using DDMC clusters (A), unclustered signaling data (B), k-means clusters (C) and GMM clusters (D). Predictive performance of a logistic regression model regressed against EGFRm/ALKf mutational status. Note that a complete portion of the entire data set containing 1311 peptides was used for every modeling strategy other than DDMC. DDMC was fit to 30561 peptides that include at least a minimum of 2 10-plex TMT experiments.
